# Deciphering acquired resistance mechanisms to sustained auxin-inducible protein degradation in cells and mice

**DOI:** 10.1101/2025.09.21.677607

**Authors:** Judith Hyle, Zhenling Liu, Shaela Fields, Jifeng Yang, Xinyan Chen, Wenjie Qi, Qiong Zhang, Byoung-Kyu Cho, Young Ah Goo, Xiuling Li, Jack Sublett, Qianqian Li, Liusheng He, Jonathon Klein, Peng Xu, Shondra M. Pruett-Miller, Beisi Xu, Chunliang Li

## Abstract

Targeted protein degradation is a favorable strategy for studying the immediate downstream effects of protein loss-of-function. An appealing platform among these technologies is the auxin-inducible degron (AID) system. Although this system has been applied extensively to cell and animal models, degradation resistance to long-term auxin treatment has not been studied. With the advent of the new AID2 system, cellular toxicity due to the high concentrations of auxin required in the original AID1 system is no longer a concern, making it possible to study protein degradation over extended periods. In this study, we derived multiple miniAID-tagged knock-in human cell lines and a Ctcf-miniAID knock-in mouse strain to investigate mechanisms of degradation resistance. We revealed four independent resistance mechanisms, including a nonsense mutation in the *CTCF* coding sequence that removed the miniAID peptide, a missense point mutation in the *miniAID* coding region that disrupted ubiquitin complex targeting, and silencing of the OsTIR1 adaptor protein. Resistance to auxin degradation was also acquired in mouse primary Ctcf^miniAID/miniAID^ knock-in B-ALL cells through missense mutations of the OsTIR1^(F74G)^ protein *in vivo* and *ex vivo*. In summary, our innovative study expands our understanding of the AID system and cautions careful consideration of design for future applications in mammalian system.

## Introduction

Due to the rapid development of genomic profiling technologies, numerous genes were discovered as essential players in human development and disease progression^1,2^. To dissect the functional roles of these genes, researchers have employed genetic loss-of-function approaches, including shRNA^3,4^, Cre/LoxP^5,6^, and recent genome-editing tools such as CRISPR/Cas9^7–9^. While these tools are powerful for impairing gene function, they also share common drawbacks. These include a longer time to complete knockdown and off-targeting effects, especially when using DNA and RNA-level editing techniques. Therefore, novel loss-of-function strategies that harness the cell intrinsic machinery to promote the degradation of the protein of interest have emerged as promising approaches to fill the gap^10–14^.

Degrons, small peptide sequences recognized by the E3 ubiquitin ligase, can be engineered to proteins of interest to allow acute protein degradation by the cellular ubiquitin-proteasome system (UPS). Common degrons include dTAG^14^, the PROTAC (proteolysis-targeting chimera) system^11,13^, Bromotag^12^, and the auxin-inducible degron (AID) system^15^. The AID system stands out because the miniAID tag is the smallest degron at only seven kDa long, minimizing the risk of altering native protein structure and function. Although first identified in plants, the AID system was introduced into mammalian cells to efficiently, rapidly, and reversibly degrade a protein of interest^16–19^. Mechanistically, the AID system conjugates the miniAID-tagged protein to the endogenous UPS through auxin (indole-3-acetic acid, IAA), facilitating specific protein degradation within a few minutes to hours^15^. To functionally equip the system in live cells, a 204-bp miniAID sequence is delivered into the endogenous locus of the targeted gene by genome editing to produce a miniAID fusion protein. In addition, the F-box adaptor protein, OsTIR1, is ectopically expressed in cells and serves as a molecular bridge, bringing together the UPS and miniAID-tagged protein in the presence of the auxin ligand. Upon auxin treatment, the ternary UPS complex immediately organizes and executes protein degradation that can be reversed upon auxin removal. Over the past decade, the generalizable application of the AID system has been reported in numerous organ systems, including worms, flies, human and mammalian cell lines, as well as the mouse model *in vivo*^15,19–21^. A few years ago, the system was upgraded to the AID2 version by generating a mutant form of OsTIR1, OsTIR1^(F74G)^, that specifically binds a synthetic auxin analog, 5-phenyl-IAA (5-Ph-IAA). The AID2 system notably decreased auxin toxicity and increased sensitivity, further optimizing it for studying immediate gene function and long-term treatment regimens *in vitro* and *in vivo*^19,22,23^.

However, as with many of the drug-based approaches, long-term drug treatment to induce degradation could lead to resistance^24,25^. Indeed, acquired resistance to PROTAC has been shown to develop through genetic alterations in key components of the UPS complex^26–28^. Because the AID system also relies on the cell’s intrinsic E3 ubiquitin ligase machinery, we hypothesized that long-term auxin treatment would enable cells to develop mechanisms to counteract protein degradation. This could lead to drug resistance, particularly for essential genes required for cell survival. To our knowledge, no systematic investigation of AID resistance has been conducted in mammalian systems, hindering our understanding of safety for future applications.

To this end, we have derived multiple human miniAID-protein knock-in cell lines against genes defined as survival essential in the acute lymphoblastic leukemia cell line SEM, such as *CTCF*, *RBM5*, and *MBNL1*. Upon long-term auxin treatment, miniAID-tagged RBM5 and MBNL1 retained auxin sensitivity. Strikingly, the miniAID-tagged CTCF frequently acquired auxin treatment resistance in human cell lines. To further explore whether the resistance phenotype is due to *in vitro* culture conditions, we confirmed the resistance phenotype from a newly established Ctcf-miniAID knock-in mouse model. Overall, in the human CTCF^AID2^ SEM cells, three independent resistance mechanisms were discovered: (1) a nonsense mutation of the *CTCF* coding sequence to truncate the miniAID peptide; (2) a missense point mutation at the *miniAID* coding region to disrupt ubiquitin complex targeting; and (3) epigenetic silencing of the OsTIR1^(F74G)^ adaptor protein. In mouse primary Ctcf^miniAID/miniAID^ knock-in B-ALL cells, missense mutations of the OsTIR1^(F74G)^ adaptor protein were acquired *ex vivo* and *in vivo*. In summary, our comprehensive study has revealed the detailed molecular mechanisms of AID resistance in mammalian and human cell systems. These findings significantly expand our understanding of the AID system and encourage careful consideration in the design of future applications.

## Results

We previously engineered the miniAID tag to an essential gene *CTCF*^29^, which is required for maintaining genome-wide chromatin architecture organization in the acute lymphoblastic leukemia B-ALL cell line SEM. CTCF^AID2^ cells have a miniAID-mClover3 tag in frame with both alleles of the 3’ end of *CTCF*. The cells exogenously express OsTIR1^(F74G)^, a mutant of OsTIR1 that binds to SKP1 to form the SKP1, CUL1, F-box protein (SCF) complex, which promotes ubiquitination and degradation of miniAID-tagged proteins in the presence of the auxin analog 5-Ph-IAA (**Fig. 1a**). In this study, we combined CTCF^AID2^ cells with extended 5-Ph-IAA treatment to determine how the system would respond to long-term CTCF depletion^23^. The previously designed CTCF^AID2^ cells, designated Clone 27, expressed OsTIR1^(F74G)^ from the lentiviral cassette pCDH-MND-OsTIR1^(F74G)^-P2A-EGFP^miniAID^ (**Fig. 1b**)^23,30^. Given that Clone 27 has been cultured *in vitro* over time and that pre-existing mutations may appear, additional CTCF^AID2^ clones were newly established by genome editing in SEM cells. Like Clone 27, the new clones included the miniAID-mClover3 tag at the 3’ end of both *CTCF* alleles, but OsTIR1^(F74G)^ was expressed from a lentiviral cassette, pCDH-MND-OsTIR1^(F74G)^-P2A-Zeocin^R^, enabling sustained Zeocin drug selection pressure to maintain OsTIR1^(F74G)^ expression (**Supplementary Fig. 1a**). These clones were designated Clone 3.2, Clone 5, Clone 17.2, Clone 20, and Clone 26 and used for the long-term auxin treatment studies (**Fig. 1b**).

**Fig. 1.**
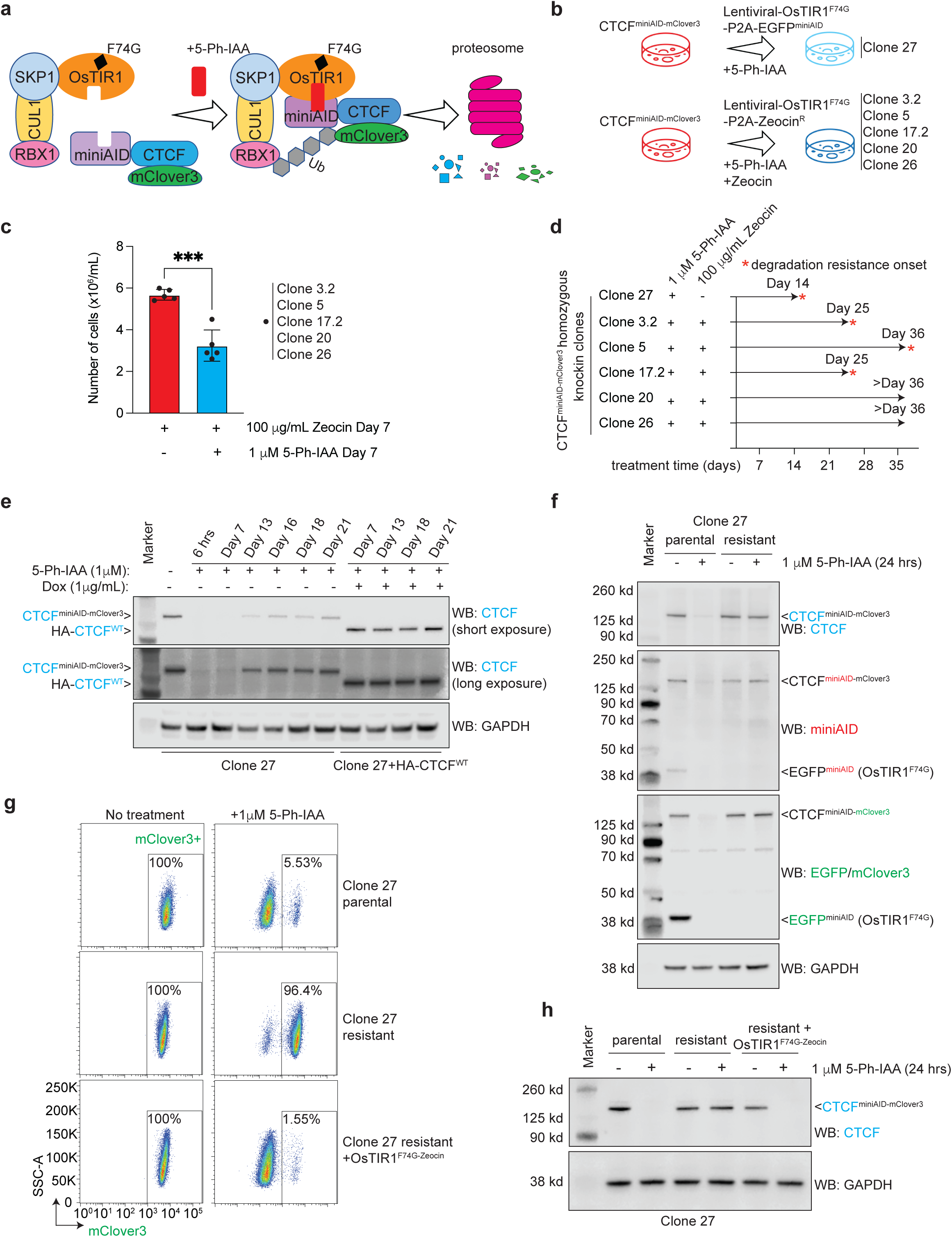
Exploring auxin-inducible degradation resistance in human cell culture. **a.** Schematic diagram detailing how the auxin-inducible system works to degrade miniAID-tagged proteins. The auxin analog, 5-Ph-IAA, acts as a ligand to bind OsTIR1^(F74G)^ to miniAID-mClover3-tagged CTCF. The SCF complex is recruited to bind OsTIR1^(F74G)^. The miniAID-mClover3-tagged CTCF is subsequently ubiquitinated by E3 ligase, and proteasomal cleavage occurs. **b.** Diagram depicting the long-term auxin culture conditions for CTCF^AID2^ clones. All clones were grown in suspension cell culture and treated with 1 μM 5-Ph-IAA for up to 36 days. Clones 3.2, 5, 17.2, 20, and 26 were also treated with 100 μg/mL Zeocin to maintain selection pressure for OsTIR1^(F74G)^ expression. **c.** Bar graph representing cell counts on day 7 in culture from CTCF^AID2^ clones treated with 100 μg/mL Zeocin only or 1 μM 5-Ph-IAA plus 100 μg/mL Zeocin. Clones were initially plated at the same cell density (1 x 10^6^/mL). The media was refreshed once on the fourth day of treatment. Cells were counted on the 7^th^ day of treatment. *** P value ≤ 0.001 calculated by unpaired *t*-test. N=5. **d.** Timeline representing the emergence of acquired auxin resistance in CTCF^AID2^ clones. **e.** Immunoblots from long-term 5-Ph-IAA culture of Clone 27 and Clone 27 + HA-CTCF^WT^. All cultures were maintained under continuous 1 μM 5-Ph-IAA treatment. Clone 27 + HA-CTCF^WT^ was also treated with 1 μg/mL doxycycline (Dox) to induce exogenous HA-CTCF^WT^. Lysates were blotted for CTCF, and GAPDH was included as a loading control. The CTCF^miniAID-mClover3^ protein can be distinguished from HA-CTCF^WT^ by its larger molecular mass. **f.** Immunoblots of Clone 27 parental and resistant cells with or without 24 hours of 1 μM 5-Ph-IAA treatment. CTCF and miniAID immunoblots confirmed that resistant cells were insensitive to auxin treatment. The GFP immunoblot showed that the OsTIR1^(F74G)^-P2A-EGFP^miniAID^ protein was no longer detected in resistant cells. GAPDH was included as a loading control. **g.** Flow cytometry analysis of mClover3 fluorescence, which corresponds to the protein expression of CTCF^miniAID-mClover3^, in Clone 27 parental cells, resistant cells, and resistant cells plus OsTIR1^(F74G)^-Zeocin^R^ with or without 1 μM 5-Ph-IAA treatment for 24 hours. Percentages of mClover3+ cells were represented in the lower-right quadrant. **h.** Immunoblot of CTCF expression in Clone 27 parental cells, resistant cells, and resistant cells plus OsTIR1^(F74G)^-Zeocin^R^ with or without 1 μM 5-Ph-IAA treatment. Restoring OsTIR1^(F74G)^ to resistant cells reestablished auxin-inducible degradation of CTCF. GAPDH was included as a loading control.

We previously established in CTCF^AID2^ cells that 1 μM 5-Ph-IAA induced complete CTCF degradation 4 hours post-treatment and for up to 96 hours without changing the media^23^. In this study, all six CTCF^AID2^ clones were kept under continuous 1 μM 5-Ph-IAA treatment for up to 36 days, with media and drugs refreshed every 72-96 hours. Clones 3.2, 5, 17.2, 20, and 26 were also treated concurrently with 100 μg/mL Zeocin to maintain selection pressure for OsTIR1^(F74G)^ expression. Because homozygous loss of *Ctcf* leads to early embryonic lethality in mice^31^, we anticipated that cells would not tolerate long-term depletion of CTCF. Indeed, we noticed a significant decrease in cell growth on day 7 in the treatment groups receiving both 5-Ph-IAA and Zeocin when compared to that of Zeocin-only groups, demonstrating that cellular fitness was affected by continued CTCF loss (**Fig. 1c**). Strikingly, four of the clones, Clones 27, 3.2, 5, and 17.2, eventually recovered through survival crisis, suggesting that those clones may acquire resistance to auxin-induced degradation of CTCF (**Fig. 1d**).

The initial long-term auxin treatment study was performed in Clone 27 and Clone 27 plus doxycycline-inducible exogenous HA-tagged WT-CTCF (Clone 27+HA-CTCF^WT^). The cells were treated with continuous 5-Ph-IAA treatment for 21 days. Doxycycline was concurrently administered to Clone 27+HA-CTCF^WT^ to induce HA-CTCF^WT^ expression. Immunoblotting for CTCF showed that CTCF^miniAID-mClover3^ initially degraded as expected, but the fusion protein expression gradually reappeared after day 13 of treatment. When HA-CTCF^WT^ was induced in parallel with treatment, CTCF^miniAID-mClover3^ remained sensitive to auxin (**Fig. 1e**). In the absence of WT-CTCF rescue, the cells adapted to maintain the essential protein CTCF by escaping auxin-induced degradation. These data highlight that CTCF dependency is the major cause of the acquired degradation resistance.

We defined Clone 27 cells that developed resistance to auxin-induced degradation of CTCF after 21 days of auxin treatment as “resistant”, and untreated cells were defined as “parental”. First, we investigated whether increasing the 5-Ph-IAA concentration would force degradation. However, even tenfold more 5-Ph-IAA did not sensitize resistant cells to auxin (**Supplementary Fig. 1b**). Therefore, the resistance mechanism likely relied on intrinsic cellular regulation. Since the lentiviral cassette carrying OsTIR1^(F74G)^ in Clone 27 also expressed EGFP^miniAID^, we next evaluated whether the degradation machinery maintained its function in resistant cells by examining how EGFP^miniAID^, a non-essential protein, responded to auxin. Parental cells showed that miniAID-tagged CTCF and EGFP degraded after 24 hours of 5-Ph-IAA treatment. However, resistant cells showed no EGFP^miniAID^ expression even in untreated cells, suggesting that the cells may have silenced the OsTIR1^(F74G)^-P2A-EGFP^miniAID^ expression (**Fig. 1f**). Real-time quantitative PCR (RT-qPCR) confirmed transcription of *OsTIR1^(F74G)^*was inhibited in resistant cells (**Supplementary Fig. 1c**). Therefore, we reasoned that additional OsTIR1^(F74G)^ would rescue the functionality of the AID system. When resistant cells were transduced with lentiviral OsTIR1^(F74G)^-P2A-Zeocin^R^, the CTCF^miniAID-mClover3^ sensitivity to auxin was completely restored. Resistant cells plus OsTIR1^(F74G)-Zeocin^ lost mClover3 fluorescence upon 5-Ph-IAA treatment, comparable to parental cells (**Fig. 1g**), and CTCF^miniAID-mClover3^ was not detectable by immunoblotting (**Fig. 1h**). Collectively, these data support that CTCF protein expression was restored under continuous auxin challenge by silencing an essential component of the degradation system.

To determine whether maintaining selection pressure for OsTIR1^(F74G)^ would affect the auxin-induced degradation of CTCF, long-term treatment of 1 μM 5-Ph-IAA and 100 μg/mL Zeocin was administered to CTCF^AID2^ Clones 3.2, 5, 17.2, 20, and 26, which exogenously expressed OsTIR1^(F74G)^-P2A-Zeocin^R^. All clones initially exhibited reduced cellular growth by day 7 of treatment (**Fig. 1c**). Clones 3.2 and 17.2 recovered from the growth restriction around day 25 (**Supplementary Fig. 1d**), followed by Clone 5. The immunoblotting analysis of Clone 5 confirmed that CTCF protein expression was restored (**Supplementary Fig. 1e**), and the mechanisms of Clones 3.2 and 17.2 auxin-resistance development will be explored in further detail. Clones 20 and 26 did not recover CTCF protein expression over the treatment time course (**Supplementary Fig. 1f, g**) and maintained a reduced growth rate in culture.

Immunoblotting of Clone 3.2 following 36 days of long-term (LT) treatment with 5-Ph-IAA and Zeocin showed that the endogenous CTCF^miniAID-mClover3^ became resistant to auxin-induced degradation. When LT-treated cells removed from treatment for 48 hours (designated as Clone 3.2 resistant) were re-challenged with 5-Ph-IAA, CTCF^miniAID-mClover3^ remained insensitive to auxin (**Fig. 2a**). Fluorescence of mClover3 was minimally abrogated upon 5-Ph-IAA treatment in resistant cells, demonstrating that the majority of the endogenous CTCF^miniAID-mClover3^ protein was resistant to auxin (**Fig. 2b**). Although the CTCF^miniAID-mClover3^ protein was detectable by immunoblotting for CTCF in LT- and resistant treated cells, it was no longer detectable by the AID antibody, suggesting that the AID epitope might be blocked (**Fig. 2a**). To further examine this observation, we conducted RNA-seq analysis in Clone 3.2 parental (untreated) and resistant cells and detected a C > T mutation that resulted in a proline-to-serine missense mutation at amino acid 23 (P23S) of the miniAID tag (**Fig. 2c, d**). To confirm that the P23S mutation could lead to auxin resistance, the AID on the lentiviral OsTIR1^(F74G)^-P2A-EGFP^miniAID^ cassette was mutated to express the P23S-AID. Then, 293T cells were transiently transfected with either the ^AID-WT^EGFP or ^AID-P23S^EGFP lentiviral construct and challenged with 5-Ph-IAA treatment for 24 hours. Cells expressing ^AID-WT^EGFP demonstrated acute loss of EGFP fluorescence after 5-Ph-IAA treatment. However, ^AID-P23S^EGFP expressing cells maintained EGFP fluorescence following treatment (**Fig. 2e**). Immunoblot analysis further confirmed that ^AID-P23S^EGFP was no longer degradable by auxin treatment. In addition, the AID antibody could no longer detect ^AID-P23S^EGFP, highlighting that the mutation disrupted the antibody-epitope interaction (**Fig. 2f**). The long-term treatment was repeated in parental Clone 3.2 to determine whether auxin resistance was reproducible. Indeed, auxin-resistant CTCF^miniAID-mClover3^ appeared around day 25 of treatment, and Sanger sequencing confirmed the P23S-AID mutation in cDNA and genomic DNA from cells treated for 34 days (**Supplementary Fig. 2a-c**). Collectively, these data support that, based on the selection pressure, CTCF^miniAID-mClover3^ acquired a genomic mutation to impair the function of miniAID and retain the expression and function of CTCF in cells.

**Fig. 2.**
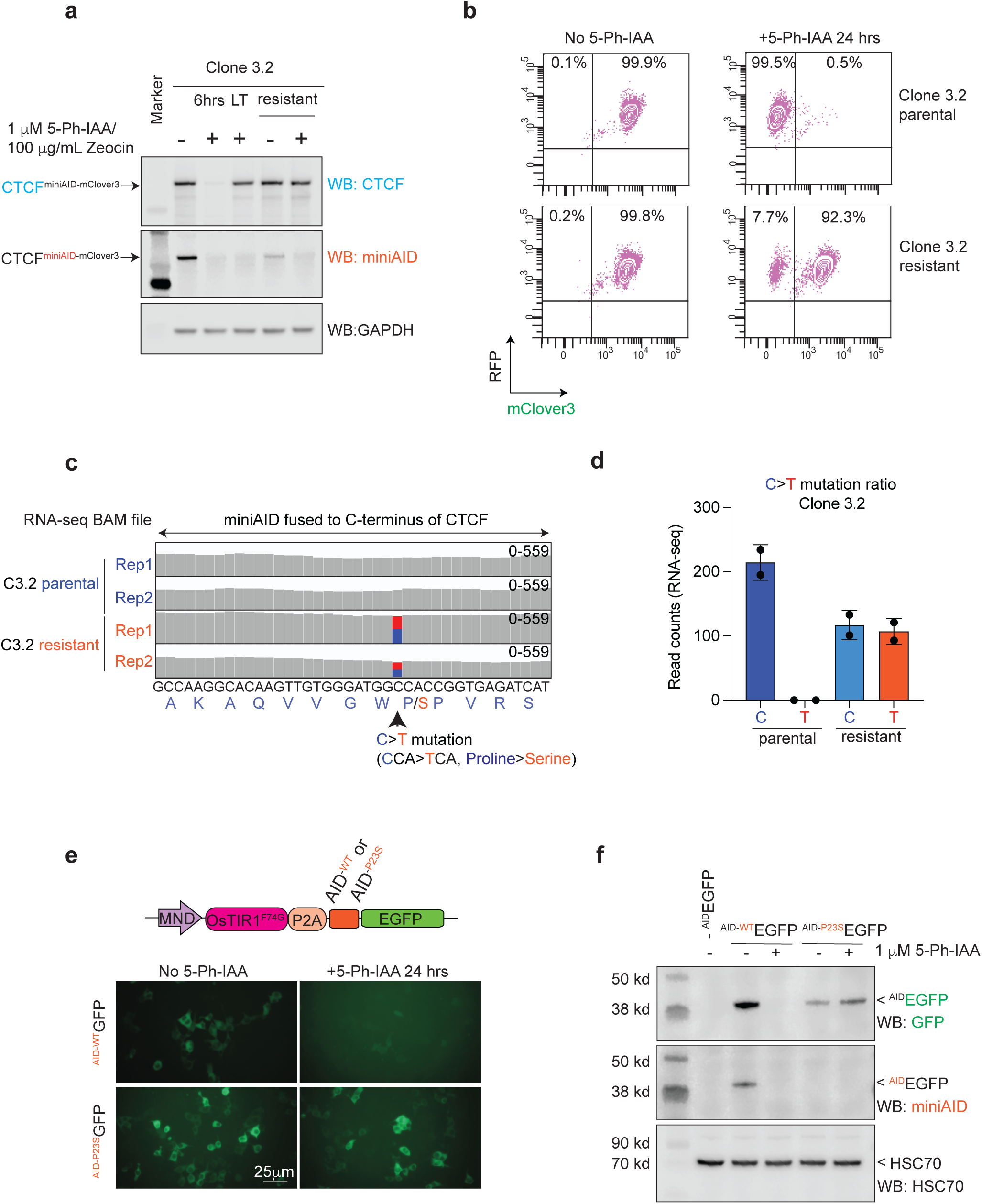
Acquired genetic mutation of degron revealed upon long-term auxin treatment. **a.** Immunoblots of lysates from Clone 3.2 with and without 1 μM 5-Ph-IAA/ 100 μg/mL Zeocin treatment for 6 hours (6 hrs) and 36 days (LT), and LT-treated cells with drugs washed out for 48 hours (resistant), also with or without 1 μM 5-Ph-IAA/ 100 μg/mL Zeocin for 24 hours. CTCF immunoblot showed that CTCF^miniAID-mClover3^ became resistant to degradation in LT and in resistant cells. However, the CTCF^miniAID-mClover3^ protein was no longer detectable by miniAID. GAPDH was included as a loading control. **b.** Flow cytometry analysis of RFP (OsTIR1^(F74G)^-P2A-Zeocin^R^-EF1α-RFP) and mClover3 (CTCF^miniAID-mClover3^) fluorescence in Clone 3.2 parental and resistant cells with or without 1 μM 5-Ph-IAA treatment for 24 hours. Percentages of RFP+ and mClover3+ cells are represented in the upper-right quadrant. RFP+ and mClover3-cells are represented in the upper-left quadrant. **c.** RNA-seq track of the miniAID cDNA sequence from RNA collected from Clone 3.2 parental and resistant cells showing the C > T mutation at amino acid 23, resulting in a proline to serine missense mutation. The blue bar represents the wild-type “C” nucleotide. The red bar represents the “T” mutation. **d.** Bar graph representing read counts of the C > T mutation ratio in Clone 3.2 parental and resistant cells. The “T” mutation was not detected in parental cells. **e.** Fluorescence microscopy of 293T cells transduced with OsTIR1^(F74G)^-P2A-EGFP^miniAID^ carrying either AID^WT^ or AID^P23S^ EGFP. AID^P23S^ EGFP maintained GFP fluorescence after 1 μM 5-Ph-IAA treatment for 24 hours in cells expressing ^AIDP23S^EGFP. **f.** Immunoblots of lysates from 293T cells transduced with OsTIR1^(F74G)^-P2A-EGFP^miniAID^ carrying either ^AIDWT^EGFP or ^AIDP23S^EGFP following 24 hours of 1 μM 5-Ph-IAA treatment. The CTCF immunoblot shows that the ^AIDP23S^EGFP protein was no longer sensitive to auxin-induced degradation, and the miniAID immunoblot reveals that the ^AIDP23S^EGFP protein was no longer detectable by the miniAID antibody. HSC70 was included as a loading control.

Unlike Clone 3.2, immunoblotting of Clone 17.2 following LT treatment (36 days) with 5-Ph-IAA and Zeocin showed that the endogenous CTCF^miniAID-mClover3^ band remained sensitive to auxin degradation. However, the emergence of a smaller truncated band was observed (**Fig. 3a**). Although the CTCF antibody detected the truncated band, it was not detectable by the AID antibody, suggesting that the protein had undergone a C-terminus truncation. Auxin and Zeocin were washed out from LT-treated cells for 48 hours and were defined as resistant. Following wash-out, resistant cells were re-challenged with 5-Ph-IAA and Zeocin for 24 hours. Immunoblotting showed that the endogenous CTCF^miniAID-mClover3^ band was restored without auxin and remained sensitive to auxin treatment. The truncated CTCF band that developed during the LT treatment was maintained following wash-out (**Fig. 3a**). Fluorescence analysis also confirmed that mClover3 fluorescence was reduced in Clone 17.2 resistant cells following auxin treatment, further demonstrating the sensitivity of endogenous CTCF^miniAID-mClover3^ to auxin-induced degradation (**Fig. 3b**). To systematically map and characterize the truncated CTCF, CTCF antibody-based immunoprecipitation and mass spectrophotometry (IP-MS) were performed on parental (untreated) and Clone 17.2 resistant cells treated with 5-Ph-IAA to remove the endogenous CTCF^miniAID-mClover3^. Clone 3.2 resistant cells were also included. While IP-MS captured peptides covering the entire sequence of CTCF^miniAID-mClover3^ in both parental and Clone 3.2 resistant cells, Clone 17.2 resistant cells displayed an abridged sequence for CTCF^miniAID-mClover3^, with a likely breakpoint occurring before exon 12 that removed the 3’ end of CTCF along with the miniAID-mClover3 tag (**Fig. 3c, d and Supplementary Fig. 3a**). RNA-seq further confirmed a C > T mutation that resulted in a nonsense mutation of glutamine to a stop codon at amino acid 666 (Q666*) in exon 11 (**Fig. 3e and Supplementary Fig. 3b**). Long-term treatment of Clone 17.2 was performed independently for two additional replicates to determine whether the truncated CTCF would stably appear upon auxin challenge. Deep sequencing confirmed that the C > T mutation emerged in conjunction with the expression of the truncated CTCF protein, CTCF^Q666*^ (**Fig. 3f and Supplementary Fig. 3c, d**). Notably, pan-cancer genome and transcriptome analysis of pediatric malignancies identified a splice region and frameshift mutation at P667 at the E11/E12 junction of *CTCF* in cases of AML and neuroblastoma, respectively, supporting that this region of the gene is predisposed to genetic instability^32^.

**Fig. 3.**
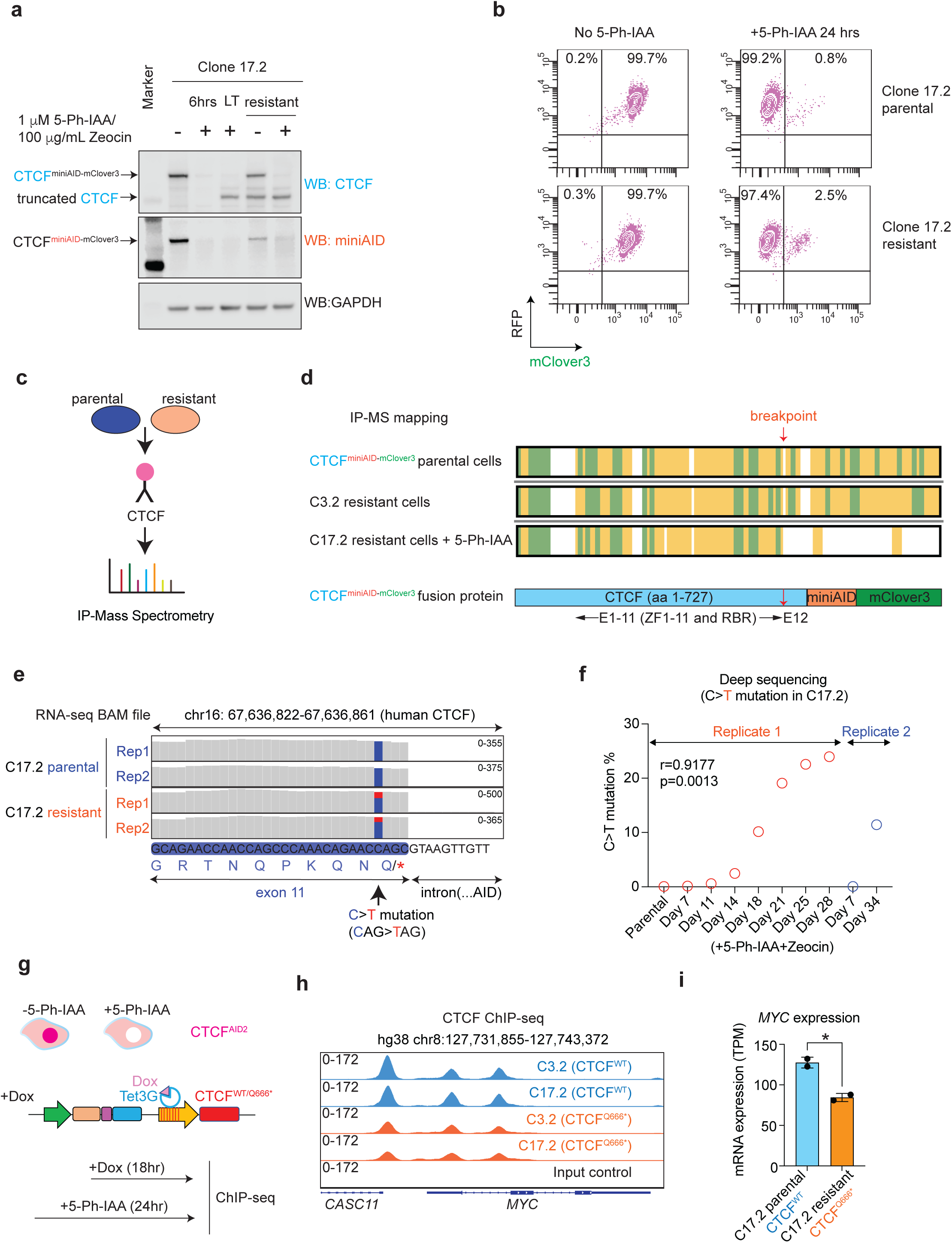
Acquired genetic mutation of CTCF revealed upon long-term auxin treatment. **a.** Immunoblots of lysates from Clone 17.2 with or without 1 μM 5-Ph-IAA/ 100 μg/mL Zeocin treatment for 6 hours (6 hrs) and 36 days (LT), and LT-treated cells with drugs washed out for 48 hours (resistant), also with or without 1 μM 5-Ph-IAA/ 100 μg/mL Zeocin for 24 hours. CTCF immunoblot showing CTCF^miniAID-mClover3^ remained sensitive to degradation in LT and resistant cells. However, a truncated CTCF appeared in LT-treated cells. MiniAID did not detect the truncated CTCF protein. GAPDH was included as a loading control. **b.** Flow cytometry analysis of RFP (OsTIR1^(F74G)^-P2A-Zeocin^R^-EF1α-RFP) and mClover3 (CTCF^miniAID-mClover3^) fluorescence in Clone 17.2 parental and resistant cells with or without 1 μM 5-Ph-IAA treatment for 24 hours. Percentages of RFP+ and mClover3+ cells are represented in the upper-right quadrant. RFP+ and mClover3-cells are represented in the upper-left quadrant. **c.** Schematic diagram of CTCF immunoprecipitation and mass spectrophotometry of parental and resistant CTCF^AID2^ clones 3.2 and 17.2. **d.** Sequence bar representing mass spectrometry peptide sequence coverage of the CTCF^miniAID-mClover3^ protein. Yellow and green shading highlight peptides detected with high confidence. Green shading predicts peptides with possible posttranslational modifications. The CTCF^miniAID-mClover3^ bar represents the CTCF amino acid sequence (blue), miniAID tag (orange), and the mClover3 tag (green). E = exon; ZF = zinc finger; RBR = RNA binding domain. The red arrow indicates the breakpoint. **e.** RNA-seq track of the exon 11 cDNA sequence from RNA collected from Clone 17.2 parental and resistant cells showing the C > T mutation at amino acid 666, resulting in a glutamine to stop codon nonsense mutation. The blue bar represents the WT “C” nucleotide. The red bar represents the “T” mutation. **f.** Chart representing the deep sequencing readout percentage of the C > T mutation’s emergence throughout continuous treatment with 1 μM 5-Ph-IAA/100 μg/mL Zeocin for up to 34 days in Clone 17.2. Two replicates were included. **g.** Schematic diagram of the model to switch from endogenous CTCF expression to doxycycline-inducible exogenous CTCF^WT-HA^ and CTCF^Q666*-HA^ expression. CTCF^AID2^ cells were transduced to express either CTCF-WT-HA or CTCF-Q666*-HA. First, cells were treated with 1 µM 5-Ph-IAA for 24 hours. After 6 hours of treatment, doxycycline (Dox) was added to induce exogenous CTCF expression. **h.** CTCF ChIP seq tracks of CTCF^AID2-WT-HA^ and CTCF^AID2-Q666*-HA^ clones (C3.2 and C17.2) at the *MYC* locus showed increased CTCF binding in CTCF^AID2-WT-HA^ clones. **i.** Reduced expression (TPM) of *MYC* observed in C17.2-resistant cells that express the truncated CTCF^Q666*^ protein when compared to C17.2 parental cells that express the full-length CTCF^WT^. P-value ≤ 0.05 calculated by unpaired *t*-test.

To determine if the truncated CTCF^Q666*^ would recapitulate CTCF^WT^ function, we compared the DNA binding affinity of CTCF^Q666*^ to CTCF^WT^ by ChIP-seq. Since the truncated CTCF arose over a long-term culture under a drug challenge, alterations to CTCF-DNA binding could not be distinguished from culture stress. Therefore, an inducible expression cell model was developed to ectopically express CTCF^WT^ or CTCF^Q666*^ upon doxycycline treatment. CTCF^AID2^ clones 5, 3.2, and 17.2 were transduced to express doxycycline-inducible CTCF^WT^ or CTCF^Q666*^. Cells were treated for 24 hours with 1µM 5-Ph-IAA to degrade endogenous CTCF. Six hours into treatment, doxycycline was added to induce exogenous CTCF expression (**Fig. 3g**). Initial immunoblots of the exogenous CTCF expression in CTCF^AID2-WT-HA^ and CTCF^AID2-Q666*-HA^ cells showed that CTCF^Q666*^ expression was less than that of CTCF^WT^. Therefore, a doxycycline titration was performed, and the dosage to induce comparable expression between CTCF^AID2-WT-HA^ and CTCF^AID2-Q666*-HA^ cells was determined to be 0.05 µg/mL and 1.0 µg/mL doxycycline, respectively (**Supplementary Fig. 4a**). CTCF ChIP-isolated the DNA regions bound to CTCF binding sites in either CTCF^WT^ or CTCF^Q666*^. Principal component analysis (PCA) confirmed that the reproducible variation among the samples was due to the mutation of CTCF (**Supplementary Fig. 4b**). Overall, 203,262 CTCF DNA binding peaks were called. Global CTCF binding was maintained in all groups, as expected, since the core zinc finger DNA binding domain remained intact in the CTCF^Q666*^ mutant. In 7,727 peaks, CTCF binding was greater in CTCF^AID2-WT-HA^ cells compared to CTCF^AID2-Q666*-HA^ cells (log2 FC > 1, FDR < 0.05). Binding was reduced at only 59 peaks in CTCF^AID2-WT-HA^ cells compared to CTCF^AID2-Q666*-HA^ cells (log2 FC > 1, FDR < 0.05) (**Supplementary Fig. 4c**). Motif enrichment analysis confirmed the CTCF binding motif as the most enriched motif among the 7,727 increased binding peaks (**Supplementary Fig. 4d**). The *MYC* locus, which is positively regulated by CTCF, showed decreased CTCF binding and transcription in CTCF^AID2-Q666*-HA^ cells (**Fig. 4h, i**). Conversely, at the *BLCAP* locus where CTCF acts as an insulator of transcription, CTCF binding increased in CTCF^AID2-Q666*-HA^ cells (**Supplementary Fig. 4e**). However, most loci showed equal CTCF binding (**Supplementary Fig. 4e**). Taken together, upon selection pressure in culture, the *CTCF^miniAID-mClover3^* acquired an alternative genomic mutation to kick off miniAID and retain the near-functional peptide fraction of CTCF at a genome-wide scale.

**Fig. 4.**
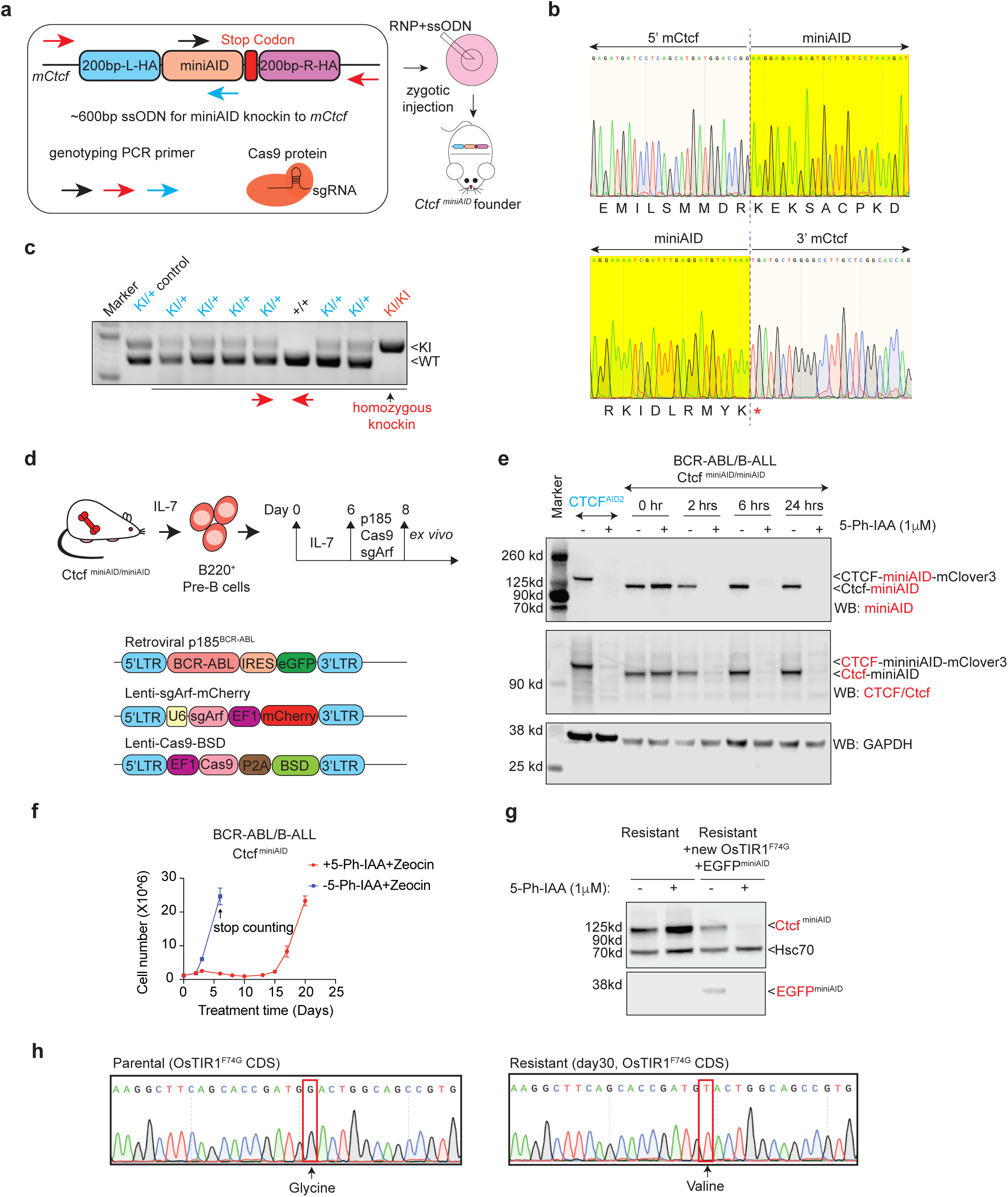
Establish and characterize the Ctcf^miniAID/miniAID^ mouse strain and Ctcf^miniAID/miniAID^ BCR-ABL B-ALL model for studying acute and long-term response to auxin-induced degradation. **a.** Schematic diagram showing the 600bp single-stranded oligodeoxynucleotides (ssODN) with 200-bp 5’ and 3’ homology arms (200bp-L-HA and 200bp-R-HA) and a 204-bp miniAID sequence. The arrows indicate primer positions for genotyping PCR. The ribonucleotide complex (RNP) is formed by the Cas9 protein and sgRNA against *Ctcf in vitro*. The ssODN was mixed with the RNP complex, followed by a zygotic injection to establish the Ctcf^miniAID^ founder mice. **b.** Sanger sequencing results confirm the junction sequence of 5’ and 3’ miniAID in the same reading frame with *Ctcf* before the stop codon. The top row shows the reflective codon readout of the 3’ end of *Ctcf* switching to the 5’ miniAID codon sequence. The second row illustrates the 3’ end of the miniAID tag with a stop codon transitioning back to the *Ctcf* endogenous sequence. **c.** Genotyping PCR results confirm homozygous Ctcf^miniAID/miniAID^ among wild-type and heterozygous offspring. Red arrows indicate that the genotyping PCR primers used sat outside the homology arms. **d.** Schematic diagram describing the procedure to derive the primary cell model from the homozygous Ctcf^miniAID/miniAID^ knock-in animal. Bone marrow samples from 8-week-old mice were collected and cultured with BCM5 and IL-7 conditional medium for 6-8 days to maintain pre-B cells, followed by transformation with retroviral-BCR-ABL and lentiviral-Cas9, as well as lentiviral-sgArf to transform pre-B and obtain the BCR-ABL B-cell acute lymphoblastic leukemia (B-ALL) model. **e.** Immunoblot of Ctcf and miniAID expression with or without 1 μM 5-Ph-IAA and 50 μg/mL Zeocin treatment for 0 hours (0hr), 2 hours (2 hrs), 6 hours (6 hrs), or 24 hours (24 hrs). CTCF^AID2^ cells with or without 5-Ph-IAA treatment were used as controls. Upon 5-Ph-IAA treatment, the Ctcf-miniAID fusion protein was acutely degraded as early as 2 hours post 5-Ph-IAA treatment. GAPDH served as the loading control. **f.** Cell counting was conducted for Ctcf^miniAID/miniAID^ BCR-ABL B-ALL cells with and without long-term treatment of 1 μM 5-Ph-IAA and 50 μg/mL Zeocin. Replication=3. **g.** Resistant cells transduced with additional lentiviral OsTIR1^(F74G)^-EGFP-miniAID restored the Ctcf^miniAID^ sensitivity to auxin treatment, shown by immunoblotting against Ctcf and EGFP (internal control). HSC70 served as the loading control. **h.** Sanger sequencing confirmed the missense mutation (G to T) in the OsTIR1^(F74G)^ CDS, encoding an amino acid change from GGA (glycine) to GTA (valine).

To further extend our study of degron resistance to other proteins, we used a similar genome editing approach (mixture of Cas9 protein, sgRNA, and knock-in donor plasmid) to deliver the miniAID tag to *RBM5* and *MBNL1*, two genes previously identified as essential in the SEM cell line^33,34^. ^HA-miniAID^RBM5 and MBNL1^HA-miniAID^ knock-in clones were derived in SEM cells, and knock-in cells were transduced with lentiviral-OsTIR1^(F74G)^-P2A-Zeocin^R^. Cells were grown in culture with concurrent treatment of 1 μM 5-Ph-IAA and 100 μg/mL Zeocin for up to 23 days. Interestingly, reduced proliferation was not observed in either cell line. Immunoblotting of the fusion proteins by protein-specific and HA-tag antibodies following LT treatment ( > 23 days) with 5-Ph-IAA and Zeocin showed that the endogenous ^HA-miniAID^RBM5 and MBNL1^HA-miniAID^ proteins remained sensitive to auxin degradation over the treatment time course (**Supplementary Fig. 5a, b**). These data suggest that acquired auxin resistance was selective and less likely to be initiated without selection pressure.

Given that all degron resistance experiments were conducted in human cell lines known to be susceptible to clonal adaptation and variation, we explored whether acquired auxin resistance would occur in mouse primary cells, which more accurately represent *in vivo* conditions. As it is impossible to deliver the miniAID tag to both alleles of the endogenous locus in mouse primary cells, we engineered a new Ctcf-miniAID knock-in mouse model that enables the derivation of primary cells from any tissue for degradation assays. To this end, gene editing was used to deliver a mixture of Cas9 protein, sgRNA against *Ctcf* near the stop codon, and a 600-bp single-stranded oligodeoxynucleotide carrying *Ctcf* homology arms and the miniAID fragment to mouse zygotes by microinjection (**Fig. 4a**). Luckily, three founder mice with successful knock-in were obtained. Sanger sequencing confirmed the seamless knock-in of miniAID in frame with *Ctcf* before the stop codon (**Fig. 4b, Supplementary Fig. 6a**). One male founder mouse was bred for germline transmission and produced viable offspring with homozygous Ctcf^miniAID/miniAID^ alleles (**Fig. 4c, Supplementary Fig. 6b**). To determine the Ctcf-miniAID fusion protein expression in the knock-in animals, liver, spleen, and kidney tissues were collected from wild-type, heterozygous, and homozygous mice, followed by immunoblotting with the Ctcf antibody. As expected, all tissues expressed the Ctcf-miniAID fusion protein with a molecular weight 7 kDa higher than that of the wild-type Ctcf protein (**Supplementary Fig. 6c**).

To derive a primary cell model from the homozygous Ctcf^miniAID/miniAID^ knock-in animal, we collected bone marrow from an 8-week-old animal to isolate pre-B cells. The pre-B cells were subsequently cultured with BCM medium supplemented with IL-7, followed by transformation with retroviral-BCR-ABL (GFP+), lentiviral-Cas9, and lentiviral-sgArf (mCherry+), leading to the establishment of the classic double-positive BCR-ABL B-cell acute lymphoblastic leukemia (B-ALL) model (**Fig. 4d, Supplementary Fig. 6d**)^35,36^. These Ctcf^miniAID/miniAID^ BCR-ABL B-ALL cells exhibited a fast proliferation rate and sensitivity to the second-generation tyrosine kinase inhibitor Dasatinib, both classic signatures of BCR-ABL transformed cells (**Supplementary Fig. 6e, f**). They also demonstrated classic features of pre-B cells, including high expression levels of B220 and negative expression of IgM (**Supplementary Fig. 6f**). The established BCR-ABL Ctcf^miniAID/miniAID^ cells were then transduced with lentiviral OsTIR1^(F74G)^-P2A-Zeocin^R^. Upon 5-Ph-IAA treatment, the Ctcf-miniAID fusion protein was acutely degraded as early as 2 hours post-treatment, and expression was restored after auxin removal (**Fig. 4e and Supplementary Fig. 6g**).

*Ctcf* was considered an essential gene for survival in almost all cell types. To confirm its role in our Ctcf^miniAID/miniAID^ BCR-ABL B-ALL cells, a competitive proliferation assay (CPA) was conducted by infecting cells with lentiviral-Cas9 and three sgRNAs against the coding exons of *Ctcf*. All guide RNAs were cloned into a lentiviral cassette with CFP fluorescence. A flow cytometry analysis tracing the CFP fluorescence profile of the cells over 8 days showed cellular fitness dependency on the positive control *Myc* and *Ctcf*, supporting *Ctcf* as being essential for survival (**Supplementary Fig. 6h, i)**. To explore whether acquired resistance during long-term Ctcf^miniAID^ degradation would arise due to the survival crisis, long-term treatment of 1 μM 5-Ph-IAA and 50 μg/mL Zeocin was administered to the cell culture. Cells initially exhibited a dramatic growth crisis immediately upon treatment and then recovered around day 14, followed by a similar expansion rate compared with growth of parental cells thereafter (**Fig. 4f**). Meanwhile, the resistant cells remained sensitive to Dasatinib (**Supplementary Fig. 7a**). Immunoblotting analysis confirmed that full-length Ctcf protein expression was restored in resistant cells (**Supplementary Fig. 7b**). Because RNA-seq analysis failed to identify any genetic mutations in the *Ctcf* or *miniAID* sequences, we hypothesized that the resistance mechanism could be due to mutations in the adaptor protein, OsTIR1^(F74G)^. Indeed, re-introducing lentiviral OsTIR1^(F74G)^-P2A-EGFP^miniAID^ to the cells re-established the AID system’s functionality (**Fig. 4g**). Sanger sequencing of *OsTIR1^(F74G^*^)^ confirmed a G142V missense mutation (**Fig. 4h**).

To determine if Ctcf^miniAID/miniAID^ BCR-ABL B-ALL cells could potentiate leukemia *in vivo*, 100,000 Ctcf^miniAID/miniAID^ BCR-ABL B-ALL cells were injected into CD1 Nude mice (**Supplementary Fig. 7c**). Flow cytometry analysis of peripheral blood showed increased peripheral leukemic load (**Supplementary Fig. 7c, d**). A moribund phenotype typical of leukemia onset was observed 2-3 weeks post-injection. Spleen from sacrificed mice showed a significant increase in weight and size compared to the negative control (**Supplementary Fig. 7e, f**). H&E staining confirmed morphological abnormalities in the spleen of sick mice vs control, and IHC staining showed abundant GFP-positive leukemia cells in the spleen samples from sick mice (**Supplementary Fig. 7g, h**). These data confirmed the *in vivo* leukemogenic potential of the Ctcf^miniAID/miniAID^ BCR-ABL B-ALL engrafted cells.

To determine if auxin resistance would occur *in vivo*, Ctcf^miniAID/miniAID^ BCR-ABL B-ALL cells that had been transduced with MND-luciferase-P2A-YFP were delivered by tail vein injection to 10-12-week-old CD1 Nude female mice. Before transplantation, the cells were challenged with 5-Ph-IAA to confirm sensitivity to auxin (**Supplementary Fig. 7i**). After leukemia cells were transplanted, the administration of daily IP injedtion of PBS or 5-Ph-IAA will be conducted from the 2^nd^ day following injection (**Fig. 5a**). Before long-term treatment, a pilot experiment was done in mice with leukemia burden to test the drug efficacy *in vivo*. To this end, blood and bone marrow cells were collected at 6 and 24 hours after the first IP injection of drug or PBS, and confirmed acute auxin sensitivity to deplete the Ctcf^miniAID/miniAID^ *in vivo* (**Fig. 5b, c**). Thereafter, the long-term treatment of 5-Ph-IAA was carried out by daily IP injection to maintain the degradation pressure. Leukemia load was monitored by daily observation and weekly luciferase imaging. In the PBS/vehicle control group, 3 out of 5 mice exhibited successful engraftment of Ctcf^miniAID/miniAID^ BCR-ABL B-ALL cells, with 2/5 expressing high luciferase levels by day 15 post-injection. One mouse from the vehicle group had to be sacrificed by this time point due to high leukemic load, and the remaining two were sacrificed by day 22 after injection. In contrast, the mice administered 5-Ph-IAA showed a notably delayed onset of luminescence. However, 4 out of 5 eventually had a significant leukemic load (**Fig. 5d**). At the endpoint, peripheral blood, bone marrow, and spleen cells were immediately collected and combined in both vehicle and treatment groups. Immunoblotting against these fresh samples confirmed that Ctcf^miniAID^ had escaped degradation in the mice treated long-term with 5-Ph-IAA (**Fig. 5e**).

**Fig. 5.**
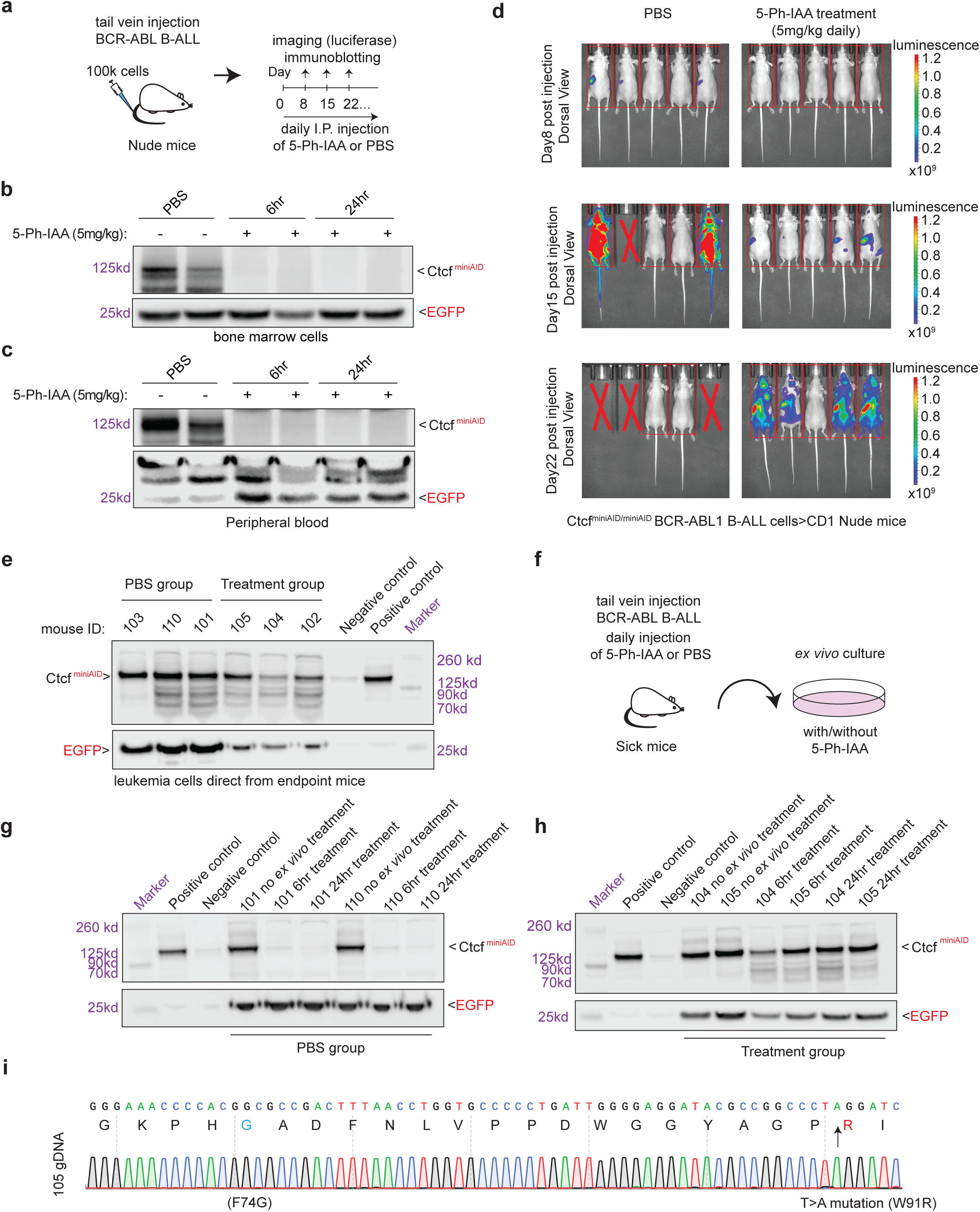
Long-term 5-Ph-IAA *in vivo* treatment induced auxin resistance. **a.** Schematic diagram summarizing *in vivo* cell injection and 5-Ph-IAA treatment. 100K Ctcf^miniAID/miniAID^ BCR-ABL B-ALL cells were injected into the Nude mice through tail-vein injection. Daily IP injection of 5-Ph-IAA (5mg/kg) or PBS was administered starting from the 2^nd^ day after cell injection. **b.** To determine how degradation efficacy *in vivo*, Ctcf^miniAID/miniAID^ BCR-ABL B-ALL cells were injected into CD1 Nude female mice, followed by a two-week expansion in host mice. Mice confirmed to have leukemia burden in their peripheral blood were then administered injections of PBS or 5-Ph-IAA. After cell injection and one dose of 5-Ph-IAA treatment, bone marrow cells were collected at 6-hour and 24-hour time points, and immunoblotting of miniAID confirmed degradation *in vivo*. EGFP was included as a loading control. **c.** After cell injection and one dose of 5-Ph-IAA treatment, peripheral blood was collected at 6-hour and 24-hour time points, and immunoblotting of miniAID confirmed degradation in vivo. EGFP was included as a loading control. **d.** Weekly luciferase imaging of the PBS vehicle group and the 5-Ph-IAA (5mg/kg) treatment group. On day 15, 3/5 mice in the vehicle group showed abundant leukemic load, while the treatment group showed minimal leukemic load. On day 22, mice in the 5-Ph-IAA (5mg/kg) treatment group demonstrated abundant leukemic load. **e.** Immunoblot analysis of miniAID and EGFP expression from the leukemia cells collected directly from endpoint mice. Both the PBS vehicle group and treatment group maintained Ctcf-miniAID fusion protein expression. **f.** Schematic diagram of *ex vivo* culture of leukemia cells from sick mice. Cells were cultured *ex vivo* and subjected to 5-Ph-IAA treatment (1μM) to confirm Ctcf degradation. **g.** Immunoblotting analysis of miniAID and EGFP expression after *ex vivo* treatment of the PBS vehicle group confirmed Ctcf degradation after 6 hours and 24 hours of 5-Ph-IAA treatment. **h.** Immunoblotting analysis of miniAID and EGFP expression after *ex vivo* treatment of the 5-Ph-IAA treatment group confirmed degradation resistance after 6-hour and 24-hour 5-Ph-IAA treatment. **I.** Sanger sequencing of the resistant cells confirmed the missense mutation (T to A) in the OsTIR1^(F74G)^ CDS.

To further determine whether the resistant phenotype could be maintained, leukemia cells collected from the peripheral blood, bone marrow, and spleen of sick mice were combined and cultured *ex vivo*. The *ex vivo* culture was challenged with 5-Ph-IAA treatment (1μM) for 6 and 24 hours (**Fig. 5f**). The vehicle group cells remained sensitive to 5-Ph-IAA treatment. In contrast, the treatment group cells exhibited resistance to auxin degradation at 6 and 24 hours post 5-Ph-IAA treatment, confirming the cells acquired auxin resistance over long-term treatment *in vivo* (**Fig. 5. g, h**). Sanger sequencing of the resistant cells confirmed the T > A missense mutation (W91R) in the OsTIR1^(F74G)^ adaptor protein (**Fig. 5i**). Strikingly, resistant cells from three individual mice carry the same point mutation. These data support that, under selection pressure, the cells acquired a genetic mutation in a crucial component of the degron system to preserve expression of the survival-essential protein, Ctcf.

In summary, we have investigated the resistance mechanisms that develop with prolonged auxin-induced degradation using knock-in cell lines and animal models. We revealed epigenetic silencing and various genetic mutations as the major mechanisms for acquired resistance (**Fig. 6**). This innovative study significantly extends our understanding of the AID system, underscoring the need for continued efforts to mitigate resistance.

**Fig. 6.**
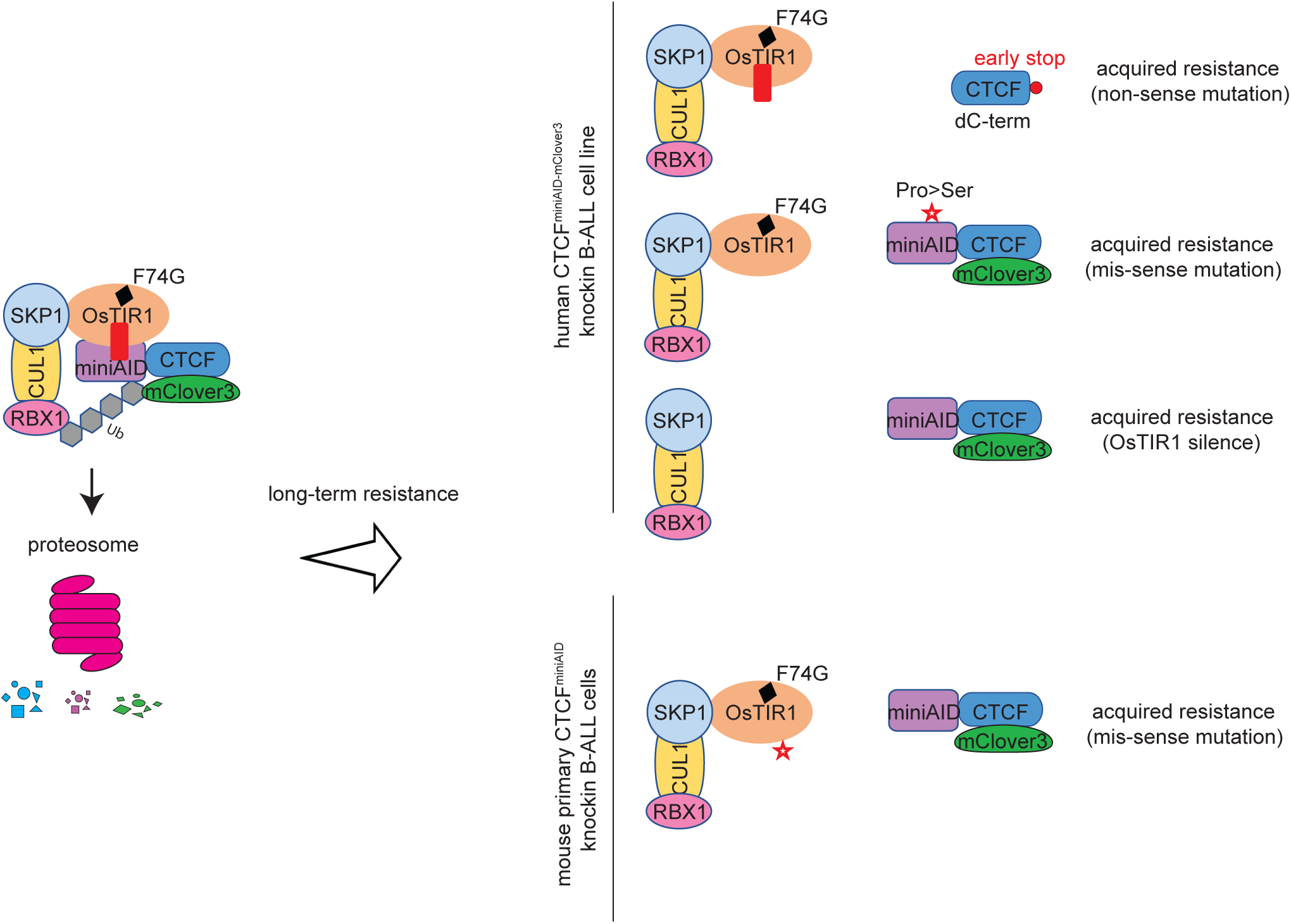
Schematic diagram summarizing the auxin-induced degron system and the auxin resistance mechanisms observed. In human CTCF^AID2^ SEM cells: (1) a nonsense mutation of the CTCF coding sequence to truncate the miniAID peptide; (2) a missense point mutation at the miniAID coding region to dissociate the E3 complex targeting; and (3) Epigenetic silencing of the OsTIR1^(F74G)^ adaptor protein. In mouse primary Ctcf^miniAID/miniAID^ knock-in B-ALL cells, a missense mutation of OsTIR1^(F74G)^ adaptor protein was acquired.

## Discussion

Targeted protein degradation has recently emerged as an appealing approach to study the immediate effects of protein loss of function. Compared to the FKBP/dTAG and PROTAC (proteolysis-targeting chimera) systems, the auxin-inducible degron (AID) is advantageous because the miniAID tag is the smallest degron at only 7kDa. Mechanistically, the AID system conjugates the SCF ubiquitination system from host cells to the AID-knock-in protein through auxin to trigger efficient, acute, and reversible protein degradation. Over the past few years, our group and others have continued to optimize and leverage this powerful tool to investigate functions of genes essential for cancer cell survival. We hypothesized that upon long-term auxin treatment, cells may evolve mechanisms to overcome protein degradation, particularly for survival-relevant genes, leading to drug resistance. To our knowledge, no investigation of AID resistance has been conducted, hindering our understanding of the safety and appropriate use of future targeted protein degradation applications.

The AID system rapidly degrades essential genes, enabling the investigation of their early roles in gene regulation and the underlying molecular mechanisms before significant cellular phenotypes emerge. Our AID model was mainly built upon the endogenous *CTCF* gene locus, a multi-functional master transcription factor that organizes chromatin architecture and plays varied roles in transcriptional regulation^29^. Homozygous knockout in mice is lethal, and complete knockout in most cells leads to impaired cell viability. Over the years, several research groups, including ours, have demonstrated the indispensable role of CTCF in maintaining genome-wide chromatin architecture, controlling chromatin accessibility, and transcription modulation^23,30,37–40^. It is well recognized that cells are addicted to CTCF expression to maintain normal physiology^31,37,41–44^. When targeting essential genes or pathways, cells hijack compensatory pathways to counter the consequences of the knockout. For instance, in high-risk MLL-rearranged leukemia cells, HOXA9-targeted cells may switch to dependency on HOXA7 and HOXA10^45^. MYC inhibition can trigger the reactivation of MYCN in many cancer cell types^46,47^. In our long-term degradation resistance study, all three mechanisms of acquired auxin resistance occurred in CTCF-miniAID models. This suggests that these cells may not have compensatory mechanisms upon CTCF loss, as it plays an essential and unique role in chromatin architecture maintenance. However, in the ^HA-miniAID^RBM5 and MBNL1^HA-miniAID^ systems, the other RBM and MBNL family members may compensate for the functional loss upon degradation. Therefore, the selective pressure is required to trigger acquired resistance to degradation. We will continue to study this hypothesis in more AID knock-in models.

Targeted protein degradation has emerged as a promising therapeutic approach to overcome the limitations of traditional therapies such as small-molecule inhibitors and chemotherapeutic agents^48^. With the advent of this technology, proteins that had previously been difficult to target became accessible for study and could be evaluated for their therapeutic potential. The PROTAC system, which utilizes a ligand to link the E3 ligase to a protein of interest, has rapidly moved to clinical trials^13^. However, the ligand binds the protein of interest through small-molecule targeting, which is not available for many proteins that are difficult to target, such as transcription factors. Additionally, resistance to PROTAC degradation has been observed with mutations arising in essential components of the E3 ligase machinery^26–28^. In this study, we have investigated the degradation resistance mechanism of a novel and complementary approach, the AID system, in human cell lines and mouse primary cells. To our understanding, this is the first and most systematic investigation to date.

The AID system utilizes the *Oryza sativa* F-box TIR1 protein, OsTIR1, instead of a linker molecule to bring the AID-tagged protein and E3 ligase machinery together. In the PROTAC system, genetic alterations to essential components of the UPS complex emerged that afforded resistance to degradation. Genetic mutations of the substrate that would abrogate ligand recognition were not seen, unlike what had previously been observed with small-molecule inhibitor resistance^49^. In contrast, acquired auxin resistance occurred through genetic alterations of *CTCF*, *miniAID*, *OsTIR1^(F74G)^*, and silencing of *OsTIR1^(F74G)^* in the AID system. Without sufficient selection pressure to enforce *OsTIR1^(F74G)^* expression, the human CTCF^AID2^ cells, Clone 27, silenced the exogenous *OsTIR1^(F74G)^* to escape targeted degradation. In addition, two separate genetic mutations were observed in CTCF^AID2^ clones that mitigated the function of the miniAID tag. These clones maintained similar RNA expression levels of the endogenous UPS complex members like *SKP1*, *CUL1*, and *RBX1* (**Supplementary Table 2**), and maintained functional OsTIR1^(F74G)^ under continuous Zeocin selection pressure. In the mouse Ctcf^miniAID/miniAID^ BCR-ABL B-ALL cells, the cells acquired mutations in the OsTIR1^(F74G^) adaptor protein in both cell culture and *in vivo* settings. Therefore, in the AID system, the cells preferred genetic alterations that inhibited the function of the degron tag or *OsTIR1^(F74G)^*over the endogenous UPS complex members. Our study emphasizes the urgent need to investigate resistance prevention before general application. For instance, the P23S (proline to serine) mutation of the AID cassette is vulnerable to genetic mutation. The AID tag could be engineered with alternative amino acids to examine whether mutation of the tag could be prevented to retain the degradation sensitivity. While the AID system has primarily been used to evaluate acute degradation of miniAID-tagged fusion proteins, advances in degron targeting, such as using single-chain AID2 antibodies (scAb-AID2) to degrade endogenous target proteins, are a step forward in the therapeutic applicability of AID technology^50^. Additional efforts are required to optimize the system for long-term degradation studies.

Recently, auxin induced resistance has appealed attentions in lower species including *Saccharomyces cerevisiaeIn*^51^ and *Yarrowia lipolytica*^52^. Our studies not only systematically characterized the degron resistance mechanism in mammalian cell system but also extended the understanding of the safety and applicability of the AID system, opening a door for others to continue optimizing it to become more powerful and generalizable.

## Methods

### Animal Model

Animal research was conducted with full ethical compliance. The protocols for mouse breeding and studies have been reviewed and approved by the St. Jude Children’s Research Hospital Institutional Animal Care and Use Committee (IACUC). All mice were maintained in sterilized conditions with a temperature of around 20 °C to 23 °C and humidity between 40% and 60%. For the miniAID knock-in mouse generation, a 600-bp single-stranded oligodeoxynucleotide (ssODN) carrying 200-bp 5’ and 3’ homology arms and a 204-bp miniAID sequence was synthesized to insert miniAID to the C-terminus of endogenous Ctcf. The ssODN (Genewiz, Inc.) was mixed with Cas9 protein (Protein Production Core Facility at St. Jude) and sgCtcf (Genewiz, Inc.), followed by a zygotic injection to get Ctcf^miniAID^ founder mice. Founder mice were bred into the C57BL/6 background for germline transmission and homozygous offspring. Genotyping primer sequences are listed in **Supplementary Table 1**.

### Vector Construction

The Lenti-Cas9-Blast vector was purchased from Addgene (Addgene #83480). The Lenti-Guide-Puro (Addgene #52963) plasmid was purchased from Addgene and subcloned to contain the IRES-CFP cassette. A pair of oligonucleotides containing a 20-bp sgRNA sequence targeting the candidate region was synthesized (Thermo Fisher Scientific) and cloned into the Lenti-Guide-Puro-IRES-CFP construct between two BsmBI sites. The retroviral expression system encoding BCR-ABL (P185) in mouse stem cell virus-internal ribosome entry site-GFP vector (MSCV-IRES-GFP) was provided by Dr. Charles J. Sherr (St. Jude Children’s Research Hospital, Memphis). The pCL20-SF2-Luc2a-YFP plasmid was provided by Dr. Martine Roussel (St. Jude Children’s Research Hospital, Memphis) and subcloned with the MND promoter after ppum1 and EcoR1 digestion to remove the CMV promoter. The pCDH-MND-OsTIR1^(F74G)^-P2A-EGFP-AID2^-^EF1α-RFP (Addgene #232800) and TRE3G-CTCF-WT-HA-MND-zeo (Addgene #232801) constructs were generated in a previous study^23^. The pCDH-MND-OsTIR1(F74G)-P2A-Zeocin^R^-EF1α-RFP construct was made by using a two-insert infusion cloning to add OsTIR1(F74G)-P2A and Zeocin^R^ to pCDH-EF1α-RFP (NEBuilder HiFi DNA assembly). The OsTIR1(F74G)-P2A cassette was amplified from pCDH-MND-OsTIR1(F74G)-P2A-EGFP-AID2-EF1α-RFP by using “Frag1 OsTIR1 F and R primers”, and the Zeocin expression cassette was amplified from TRE3G-CTCF-WT-HA-MND-zeo by using “Frag2 Zeo F and R primers”. The CMV promoter was removed by SnaB1/Xba1 restriction digestion and replaced by infusion cloning of the MND promoter amplified from TRE3G-CTCF-WT-HA-MND-Zeocin^R^ using the “pCDH MND inf F and R primers”. The pCDH-MND-OsTIR1(F74G)-P2A-Zeocin^R^ cassette was made by excising EF1α-RFP from pCDH-MND-OsTIR1(F74G)-P2A-Zeocin^R^-EF1α-RFP by restriction digestion and blunt-end cloning to circularize the plasmid. TRE3G-CTCF-WT-HA-MND-BSD was made by amplifying CTCF-WT-HA from TRE3G-CTCF-WT-HA-MND-Zeocin^R^ (Addgene #232801) and cloning by infusion into TRE3G-mCherry-MND-BSD cut with BsIW1 and EcoR1. TRE3G-CTCF-Q666*-HA-MND-BSD was made by amplifying the truncated CTCF and HA-NLS from the TRE3G-CTCF-WT-HA-MND-Zeocin^R^ plasmid and cloning by infusion into TRE3G-mCherry-MND-BSD cut with BsIW1 and EcoR1.Primers for cloning were designed with Snapgene software. All primer and plasmid sequences are listed in **Supplementary Table 1**. PCR was performed by using CloneAmp polymerase (Clontech) according to the manufacturer’s protocol, and infusion cloning was performed according to the NEBuilder HiFi DNA assembly protocol. Site-directed mutagenesis was performed to mutate the AID tag of pCDH-MND-OSTIR1^(F74G)^-P2A-EGFP-AID2-EF1α-RFP to generate pCDH-MND-OSTIR1^(F74G)^-P2A-EGFP-P23S-AID2-EF1α-RFP. AID P23S SDM F and R primers were used to amplify pCDH-MND-OSTIR1^(F74G)^-P2A-EGFP-AID2-EF1α-RFP. CloneAmp polymerase was used for the PCR, and the cycling conditions were: 98 °C for 20 seconds one cycle; 98 °C for 20 seconds, 55 °C for 20 seconds, 72 °C for one minute for 18 cycles; 72 °C for five minutes for one cycle. *Dpn1* was added to the PCR reaction to degrade the methylated DNA template. One microliter of the reaction was transformed into One Shot Stbl3 cells (Thermo Fisher Scientific). The P23S-AID clones were confirmed by Sanger sequencing.

### Generation of the AID2 Knock-in Cell Lines

The CTCF^AID2^ cell line Clone 27 was created by infecting the pCDH-MND-OSTIR1^(F74G)^-P2A-EGFP^AID2^-EF1α RFP construct into a previously derived SEM B-ALL cell line expressing the endogenous CTCF^miniAID-mClover3^ fusion protein and doxycycline-inducible wild-type OSTIR1^WT23^. The Clone 27 +HA-CTCF^WT^ was made by transducing Clone 27 with TRE3G-CTCF-WT-HA-MND-Zeocin^R^. The CTCF^AID2^ Clones 3.2, 5, 17.2, 20, and 26 were made by using a single guide RNA (sgRNA) designed to target the 3’ end of *CTCF* (Synthego), purified Cas9 protein (Protein Production Core Facility at St. Jude), and the previously described CTCF-miniAID-mClover3 donor knock-in vector to deliver a miniAID-mClover3 tag to the 3’ end of endogenous *CTCF*. The Lonza Nucleofector 4D SF kit and program EH100 were used to deliver 100 μM sgCTCF, 20 μM Cas9 protein, and 100 ng of the CTCF-miniAID-mClover3 donor knock-in vector to 50,000 SEM wild-type cells. Cells were allowed to recover before sorting for mClover3 expression. A bulk population of mClover3+ cells was transduced with pCDH-MND-OSTIR1^(F74G)^-P2A-Zeocin^R^-EF1α-RFP. After recovery, cells were sorted for double-positive mClover3 and RFP populations. This population was again sorted for single cells, and clones were screened by PCR and immunoblotting to confirm the biallelic miniAID-mClover3 knock-in. A similar knock-in strategy delivered the miniAID tag to the N-terminus of *RBM5* and the C-terminus of *MBNL1* in SEM cells. Knock-in cassettes were made by Twist. Guide RNA sequences can be found in **Supplementary Table 1**.

### Mouse Bone Marrow Cell Isolation and B-ALL Cell Culture

Mouse bone marrow cells were isolated from the long bones of 8-week-old Ctcf^miniAID/miniAID^ knock-in mice in a C57BL/6 background. Red blood cells were removed with red blood cell lysis buffer (Sigma, Cat. No. R7757). Cells were suspended and cultured in complete BCM5 and IL-7 conditional medium for 6-8 days to maintain pre-B cell expansion. Then, the cell medium was switched to the Liquid B Cell Medium (LBCM). About 100,000 cells were infected with MSCV-BCR-ABL(P185)-IRES-GFP, Lenti-Cas9-Blast and Lenti-sgArf-mCherry. A detailed recipe for the culture medium is provided in **Supplementary Table 1**. For long-term treatment, mouse primary BCR-ABL Ctcf^miniAID/miniAID^ cells were first transduced with pCDH-MND-OsTIR1^(F74G)^-P2A-Zeocin^R^. Cells were cultured in a 6-well plate with 1.2 million cells for three replicates in complete liquid B cell medium with Zeocin (50 μg/mL) and with or without 1 μM 5-Ph-IAA treatment; Medium was changed every 2-3 days. Cell numbers were counted every 2-3 days using a Countess II automated cell counter (Thermo Fisher Scientific) until the counts reached over 20 million (day 6-7 for non-5-Ph-IAA treatment, day 20 for 5-Ph-IAA treatment groups).

### Cell Line Culture

The SEM B-ALL cells were cultured in RPMI-1640 medium (Lonza) containing 10 % fetal bovine serum (FBS) (Hyclone), 2 mM glutamine (Sigma), and 1% penicillin/streptomycin (Thermo Fisher Scientific) in a 37 °C incubator with a 5 % CO_2_ atmosphere and 95 % humidity. Cells were routinely tested for mycoplasma infection (Lookout Mycoplasma PCR Detection Kit, Sigma), and the cell identity of SEM was confirmed by short tandem repeat (STR) analysis. 293T cells were cultured in DMEM medium (Gibco) containing 10 % fetal bovine serum (FBS) (Hyclone), 2 mM glutamine (Sigma), and 1% penicillin/streptomycin (Thermo Fisher Scientific) in a 37 °C incubator with a 5 % CO_2_ atmosphere and 95 % humidity. For long-term auxin treatment, Clone 27 was treated with 1 μM 5-Ph-IAA (MedChemExpress) for 21 days, with media refreshed every 72-96 hours. Clone 27 + HA-CTCF^WT^ was treated with 1 μM 5-Ph-IAA and 1 μg/mL doxycycline for 21 days, with media refreshed every 72-96 hours. Clones 3.2, 5, 17.2, 20, and 26 were initially plated at the same density, and cell counts were taken using a Countess II automated cell counter (Thermo Fisher Scientific). These clones were treated with 1 μM 5-Ph-IAA and 100 μg/mL Zeocin (Thermo Fisher Scientific) for up to 36 days, with media refreshed every 72-96 hours. ^HA-miniAID^RBM5 and MBNL1^HA-miniAID^ clones were treated with 1 μM 5-Ph-IAA and 100 μg/mL Zeocin for up to 23 days, with media refreshed every 72-96 hours.

### *In vivo* cell injection and treatment

Ctcf^miniAID/miniAID^ BCR-ABL B-ALL cells were infected with Lenti-MND-luciferase-P2A-YFP to select for a triple-positive population (GFP+, mCherry+, and YFP+). One hundred thousand cells were resuspended in 0.15 mL of PBS and implanted through tail vein injection into 10-12-week-old CD1 Nude female mice (Jackson Laboratory, stock # 005557). PBS or 5-Ph-IAA (MedChemExpress,5mg/kg body weight) was administered daily from the second day after injection. Daily observation and weekly bioluminescence imaging were applied to monitor leukemia progression. Mice were euthanized when moribund: (dehydration, ruffled fur, poor mobility, respiratory distress, > 20% body weight loss) or showed > 80% leukemia cell burden in peripheral blood. At the endpoint, peripheral blood, bone marrow cells, and spleen were collected for immunoblotting.

### MTT Assay

Cell viability was measured with 3-(4,5-dimethyl-2-thiazolyl)-2,5-diphenyl-2H-tetrazolium bromide (Sigma M5655-1G). In brief, live cells were cultured in 96-well plates with regular medium (BCR-ABL cells: 24,000 cells/well; SEM cells: 24,000 cells/well; K562: 10,000 cells/well), with a diluted concentration spectrum of Dasatinib (S1021, Selleckchem) from 10 μM to 0.001 μM for 3 days. MTT (Sigma, M5655-1G) was added to each well at 5 mg/mL. Cells were incubated at 37°C for 4 hours, followed by adding 100 μL of isopropanol/acetic acid (1 mL HCl in 250 mL isopropanol) to stop the reaction. The absorbance was measured by a microplate reader (BioTek Synergy2 plate reader) at 570 nm. The dose-response curves and IC50 values were calculated using Prism 10 software.

### Flow Cytometry

Suspension-cultured cells were collected and filtered through a 70-micron cell strainer before flow cytometry sorting for RFP, mCherry, and/or mClover3-positive cells. The same FL1/FITC channel as GFP was used to detect fluorescence from mClover3. Dead cells were excluded by DAPI. For the staining of BCR-ABL B-ALL cells, cells were mixed with 100 μL APC-B220 and PE-IgM antibody buffer (0.1% BSA PBS+ 1:50 antibody), followed by incubation in the dark room at room temperature for 20 mins. Cells were washed twice with 4mL of 0.1% BSA PBS and resuspended in 200 μL of 0.1% BSA PBS for flow analysis. Data analysis and presentation were produced by FlowJo software.

### Competitive proliferation assay

BCR-ABL pre-B cells were transduced with Lenti-Cas9-Blast, selected by blasticidin, and then transduced with individual sgRNAs to *Myc* and *Ctcf* in CFP-expressing lentiviral vectors. The day following transduction, the cells were washed three times in PBS to eliminate the virus. On the second day after transduction, the percentage of CFP-positive cells was measured by flow cytometry and set as the starting time point reference. CFP fluorescence was measured again on days 5 and 8 post-transduction. The percentage of CFP-positive cells was normalized to that at the starting time point.

### Immunoblotting

Cells were lysed in RIPA buffer, and lysates were run on an SDS-PAGE gel (Thermo Fisher Scientific), followed by protein transfer to a PVDF membrane (Bio-Rad) at 100 V for 1 hour. Membranes were blocked with 5 % non-fat milk in TBS-T (10 mM Tris, pH 8.0, 150 mM NaCl, 0.5 % Tween-20) for 1 hour at room temperature, then incubated overnight with primary antibodies at 4 °C with gentle rocking (GAPDH, ThermoFisher Scientific, AM4300; CTCF, Diagenode, C15410210-50; CTCF, Millipore, 07729; miniAID, MBL, M214-3; GFP, Santa Cruz, sc-9996; HSC70, Santa Cruz, sc-7298; HA, Cell Signaling, 3724S). Membranes were washed in TBS-T and then incubated with secondary antibodies in 5% non-fat milk/TBS-T for 1 hour at room temperature [donkey anti-rabbit IgG HRP (GE Healthcare) for CTCF and HA primary blots; sheep anti-mouse IgG HRP (GE Healthcare) for GAPDH, miniAID, GFP, and HSC70 primary blots]. After washing in TBS-T, blots were developed with ECL (Perkin Elmer) and visualized by using the Licor System.

### RNA-seq

The Zymo RNA Clean and Concentrator Kit with in-column DNase treatment (Zymo Research) was used to isolate RNA from cells. The Kapa RNA HyperPrep Kit with RiboErase (HMR) was used to prepare cDNA libraries.

### Quantitative Real-time Q-PCR

The High-Capacity cDNA Reverse Transcriptase kit (Applied Biosystems) was used to make cDNA, and the FAST SYBR Green Master Mix was used for real-time RT-qPCR (Applied Biosystems). RT-qPCR primers for GAPDH and OsTIR1^(F74G)^ are listed in **Supplementary Table 1**. The ^ΔΔ^CT method was used to determine relative expression levels^53^.

### *In vivo* bioluminescence imaging

The imaging was performed using IVIS Spectrum or IVIS-200 imaging systems. Five to ten minutes prior to imaging, animals were injected (IP) with D-Luciferin (15 mg/ml in sterile saline) at a dose of 150 mg/kg (10 µL/g of body weight). After administration, animals were anesthetized using Isoflurane and maintained via nosecone on a heated imaging bed within the system for the duration of the scan. Following imaging, animals will be allowed to recover on a heating blanket under observation and supplemented with oxygen as required.

### Amplicon-based Deep Sequencing and Sanger Sequencing

RNA was extracted from Clone 17.2 cells at various time points over the long-term auxin/Zeocin treatment, and cDNA was made. First-round PCR primers were designed to amplify an ∼ 340-bp region containing the C > T mutation in exon 11 and add the Nextera Read1 and Read2 adapter sequences. PCR products were gel-extracted and amplified in a second round of PCR with indexing primers (Illumina i5(S)/i7(N) primers). Purified indexed PCR fragments were submitted for amplicon sequencing, 150-bp single-end reads on the Illumina NovaSeq 6000. The C > T mutation of the miniAID in Clone 3.2 was amplified and sequenced by Sanger sequencing. Primer sequences can be found in **Supplementary Table 1**.

### Immunoprecipitation and Mass Spectrometry

For IP-MS, 100 million cells of each of the following conditions were collected: Clone 3.2 parental, Clone 3.2 parental + 1 μM 5-Ph-IAA for 24 hours, Clone 3.2 resistant, and Clone 17.2 resistant + 1 μM 5-Ph-IAA for 24 hours. Cells were washed once in cold PBS and then resuspended in 10 mL cold non-denaturing IP buffer (200 mM Tris pH 7.4, 137 mM NaCl, 2 mM EDTA, 1% NP-40, protease inhibitors) and rotated at 4 °C for 30 minutes. Cells were briefly sonicated, and cell lysis was monitored by trypan blue staining. Cell debris was removed from the lysates by centrifugation at 4000 rpm for 10 minutes. Lysates were transferred to a new 15 mL conical tube. CTCF antibody (Diagenode) was added at a concentration of 5 μg/mL and incubated with lysates overnight on a rotator at 4 °C. The following day, 250 μL prewashed Protein G Dynabeads (Pierce) were added to the lysates and incubated for 6 hours on a rotator at 4 °C. Tubes were placed on a magnetic stand, and lysates were removed. Beads were washed twice with IP buffer, followed by 5 washes with PBS. On the last wash, the beads were transferred to a new tube. Beads were shipped on dry ice to the Mass Spectrometry Technology Access Center at the McDonnell Genome Institute (MTAC@MGI) at Washington University School of Medicine for further analysis.

### ChIP-Seq

CTCF^AID2^ clones 5, 3.2, and 17.2 were transduced with either TRE3G-CTCF-WT-HA-BSD (CTCF^AID2-WT-HA^ clones) or TRE3G-CTCF-Q666*-HA-BSD (CTCF^AID2-Q666*-HA^ clones) to make three biological replicates for each CTCF variant. All clones were treated with 1 µM 5-Ph-IAA for 24 hours. Six hours after treatment initiation, 0.05 µg/mL or 1.0 µg/mL doxycycline was added to CTCF^AID2-WT-HA^ clones and CTCF^AID2-Q666*-HA^ clones, respectively. Twenty million cells of each were collected and fixed with 1% formaldehyde for 8 minutes at room temperature in Covaris fixation buffer (Covaris TruChIP Chromatin Shearing Kit, 520154). Chromatin was prepared according to the truChIP Chromatin Shearing Kit protocol. Chromatin was sheared on the Covaris M220 ultrasonicator set at a duty factor of 10 and 200 cycles/burst for 10 min at a set point of 6 °C. Sheared chromatin was centrifuged for 10 min at 8000 × g. Clarified chromatin was amended to a final concentration of 50 mM Tris-HCl, pH 7.4, 100 mM NaCl, 1 mM EDTA, 1% NP-40, 0.1% SDS, and 0.5% Na deoxycholate plus protease inhibitors (PI). Active Motif spike-in chromatin (53083) and antibody (61686) were added to sheared chromatin according to the manufacturer’s protocol. Ten µgs of CTCF antibody (Diagenode, c15410210-50) was also added, and the chromatin was rotated at 4°C overnight. The following day, Protein G Dynabeads (Invitrogen, 10004D) were added, and the samples were rotated at 4°C for 4 hours. Samples were placed on a magnet, and the unbound fraction was removed. Beads were washed two times in high-salt wash buffer 1 (50 mM Tris–HCL pH 7.4, 1 M NaCl, 1 mM EDTA, 1% NP-40, 0.1% SDS, 0.5% Na deoxycholate plus PI) and 1 time with wash buffer 2 (20 mM Tris–HCL pH 7.4, 10 mM MgCl_2_, 0.2% Tween-20 plus PI). For the final wash, the beads were resuspended in wash buffer 2 and transferred to a new 1.5-mL Eppendorf tube. DNA was eluted and de-crosslinked in 1X TE plus 1% SDS, proteinase K, and 400 mM NaCl at 65 °C for 4 h. Phenol, chloroform, and isopropyl alcohol were used to precipitate the DNA. Libraries were constructed using the NEBNext Ultra II NEB Library Prep Kit and NEBNext Multiplex oligos for Illumina.

### RNA-seq Data Analysis

We performed the paired-end 101-cycle sequencing on the NovaSeq 6000 sequencer (Illumina) and analyzed the data using a standard pipeline. Briefly, raw reads were trimmed using TrimGalore (v0.6.3, “--paired --retain_unpaired”) and aligned to the *Homo sapiens* reference genome GRCh37.p13(hg19) by using STAR (v2.7.9a)^54^. Gene-level read quantification was performed using RSEM (v1.3.1) on the Gencode annotation v19^55^. Differential gene expression analysis was conducted by using the TMM normalization method (genes with CPM < 1 in all samples were removed), followed by Limma-voom analysis using the “voom”, “lmFit”, and “eBayes” functions from the limma R package^56^. Complete gene expression profiling was shown in **Supplementary Table 2**.

### Mass Spectrometry and Data Analysis

Trypsin (800 ng) was added to immunoprecipitated proteins on the beads for 6 hours at 37°C. The initial digested samples were centrifuged for 2 minutes at 5,000 x g, and the supernatants were collected into fresh tubes. Beads were washed twice with 100 mM ammonium bicarbonate, and the supernatants were pooled. The resulting samples were reduced with 20 mM dithiothreitol at 37°C for 1 hour, and cysteine was alkylated with 80 mM iodoacetamide for 45 minutes in the dark. Samples were treated with 600 ng of trypsin for overnight incubation at 37°C. The resulting peptides were desalted using solid-phase extraction on a C18 Spin column and eluted with 0.1% FA in 80% ACN. Peptides were analyzed via LC-MS/MS by using a Vanquish Neo UHPLC System coupled to an Orbitrap Eclipse Tribrid Mass Spectrometer with FAIMS Pro Duo interface (Thermo Fisher Scientific). The sample was loaded on a Neo trap cartridge coupled with an analytical column (75 µm ID x 50 cm PepMap^TM^ Neo C18, 2 µm). Samples were separated using a linear gradient of solvent A (0.1% formic acid in water) and solvent B (0.1% formic acid in ACN) over 120 minutes. For MS acquisition, FAIMS switched between CVs of −35 V and −65 V with cycle times of 1.5 s per CV. MS1 spectra were acquired at 120,000 resolutions with a scan range from 375 to 1500 m/z, AGC target set at 300%, and maximum injection time set at Auto mode. Precursors were filtered by using monoisotopic peak determination set to peptide, charge state 2 to 7, dynamic exclusion of 60 s with ±10 ppm tolerance. For the MS2 analysis, the isolated ions were fragmented by assisted higher-energy collisional dissociation (HCD) at 30% and acquired in an ion trap. The AGC and maximum IT were Standard and Dynamic modes, respectively. Data were searched by using Mascot (v.3.1 Matrix Science) against a customized database consisting of the Swiss-Prot human database plus 11 isoforms of the transcriptional repressor CTCF. Trypsin was selected as the enzyme, and the maximum number of missed cleavages was set to 3. The precursor mass tolerance was set to 10 ppm, and the fragment mass tolerance was set to 0.6 Da for the MS2 spectra. The carbamidomethylated cysteine was set as a static modification, and dynamic modifications were set as oxidized methionine, deaminated asparagine/glutamine, and protein N-terminal acetylation. The search results were validated with 1% false-discovery rate (FDR) of protein threshold and 90% of peptide threshold by using Scaffold (v5.3.0 Proteome Software).

### ChIP-Seq Data Analysis

Sequencing was performed by single-end 51-cycle sequencing on the NovaSeq 6000 sequencer (Illumina). Raw reads were trimmed using TrimGalore (v0.6.3) and aligned to the Homo sapiens reference genome GRCh37.p13(hg38) genome using BWA (v0.7.17-r1198). Duplicated and low mapping quality reads were removed using “bamsormadup” function from the biobambam2 tool (v2.0.87) and samtools (version 1.9, parameter “-q 1 -F 1024”)^57^. The fragment size in each sample was estimated based on the cross-correlation profile calculated from SPP (v1.11). Fragments were extended to fragment size and normalized to 15 million reads to generate bigwig files. Macs2 was used to call peaks using parameters “-g hs --nomodel –extsize <SPP_fragmentSize>=. To identify differential peaks, the reference peaks used for comparisons were generated as follows: For each sample, both “high confidence peaks” (parameter “-q 0.05”) and “low confidence peaks” (parameter “-q 0.5”) were called. Reproducible peaks (called as a high confidence peak in at least one replicate that also overlapped with a low confidence peak in the other replicates). Differential peaks were identified using the empirical Bayes method from the limma R package. For downstream analyses, both heatmaps and Spearman’s correlation were generated by deepTool^58^.

## Data Availability

RNA-seq data generated in this study were deposited at NCBI GEO as GSE307188 (reviewer token: yrwviuquzdkhzsl). The mass spectrometry data generated in this study were deposited at PXD. The mass spectrometry proteomics data have been deposited to the ProteomeXchange Consortium via the PRIDE partner repository with the dataset identifier PXD060834 (reviewer token: ipsPHDCv3fdo). **Supplementary Tables 2 and 3** summarize the RNA-expression and ChIP-seq profiling. Code repositories collected at https://doi.org/10.6084/m9.figshare.c.6186670 included RNA-seq and ChIP-seq (https://doi.org/10.6084/m9.figshare.24803238). Raw data related to figures have been deposited in **Supplementary Table 4**, and raw immunoblots have been deposited as a “raw data file”.

## Author Contribution

Conceptualization: JH, ZL, and CL; Methodology: JH, ZL, SF, JY, XC, WQ, BC, YG, XL, JS, QZ, LH, JK, BX, PX, and SM; Investigation: JH, ZL, SF, and JY; Software and formal analysis: WQ and BX; Writing, Reviewing, and Editing: JH, ZL, and CL; Supervision, project administration, and funding acquisition: CL.

## Conflict of Interest

Dr. Chunliang Li is currently an editorial board member of *Genome Biology*. All other authors claimed no conflicts of interest.

## Acknowledgment

A former intern student from Rhodes College, Shelby Mryncza, contributed significantly to collecting preliminary data under the direct supervision of Judith Hyle. We thank the members of the Li lab for the insightful discussion and comments. We thank Drs. Ruopeng Feng and Changsheng Du for valuable discussions. We appreciate the scientific editing assistance from Dr. Cherise Guess. We gratefully acknowledge the staff of the Hartwell Sequencing Core Facility, Flow Cytometry and Cell Sorting Shared Resource, Vector Development and Production Laboratory, Center for Applied Bioinformatics, Genetically Engineered Mouse Models Shared Resource, and Center for Advanced Genome Engineering at St. Jude Children’s Research Hospital, within the Comprehensive Cancer Center of St. Jude Children’s Research Hospital. This work was funded by an American Cancer Society Scholar Grant (RSG DMC-135487, to CL), V Foundation Scholar Grant (V2021-010, to CL), and American Lebanese Syrian Associated Charities (ALSAC, to CL). Mass spectrometry analyses were performed by the Mass Spectrometry Technology Access Center at the McDonnell Genome Institute (MTAC@MGI) at Washington University School of Medicine, supported by the Diabetes Research Center/NIH grant P30 DK020579, Institute of Clinical and Translational Sciences/NCATS CTSA award UL1 TR002345, and Siteman Cancer Center/NCI CCSG grant P30 CA091842. The content of this study is solely the responsibility of the authors and does not necessarily represent the official views of the National Institutes of Health.

## Supplementary Figure Legends

**Supplementary Fig. 1 Determine auxin-inducible degradation resistance in human CTCF-miniAID knock-in cell lines.**

**a.** Schematic illustration of the Lentiviral OsTIR1^(F74G)^ cassette, pCDH-MND-OsTIR1^(F74G)^-P2A-Zeocin^R^, and the miniAID-mClover3 tag knock-in design at the *CTCF* locus.

**b.** Immunoblot of Clone 27 parental and resistant cells with a titration of 5-Ph-IAA treatment for 24 hours. The CTCF immunoblot confirms that resistant cells were insensitive to auxin treatment even at higher doses. GAPDH was included as a loading control.

**c.** Quantitative PCR showing *OsTIR1^(F74G)^* expression in Clone 27 resistant cells is lower than that of parental cells. The expression shown is relative to GAPDH. *** p-value ≤ 0.001 calculated by unpaired *t*-test. N=3

**d.** Graph depicting increasing cell counts for Clones 3.2 and 17.2 after 25 days in culture with continuous 1 μM 5-Ph-IAA/ 100μg/mL Zeocin treatment. Cell counts taken from days 25 to 34 are shown.

**e, f, g.** Immunoblots from long-term 5-Ph-IAA culture of Clones 5, 20, and 26. All cultures were kept under continuous 1 μM 5-Ph-IAA/ 100 μg/mL Zeocin treatment. No T=no treatment. Lysates were blotted for CTCF, and GAPDH was included as a loading control. CTCF^miniAIDmCLover3^ expression did not escape auxin treatment in Clones 20 and 26.

**Supplementary Fig. 2 Time-dependent tracing of miniAID mutation upon long-term auxin treatment.**

**a.** Immunoblots from long-term 5-Ph-IAA culture of Clone 3.2 kept under continuous 1 μM 5-Ph-IAA/ 100 μg/mL Zeocin. Lysates were probed for CTCF and miniAID antibody, and GAPDH was included as a loading control. No T=no treatment. The auxin-resistant CTCF appeared around day 25 of treatment. Auxin-resistant CTCF, which carried the P23S-AID mutation, was not detectable by immunoblotting with a miniAID antibody.

**b and c.** Sanger sequencing trace images from PCRs of Clone 3.2 parental and treated cells [cDNA (b) and genomic DNA (c)] of the miniAID sequence showed the C > T mutation in the miniAID tag of Clone 3.2 cells treated with 1 μM 5-Ph-IAA/ 100 μg/mL Zeocin for 34 days.

**Supplementary Fig. 3 Time-dependent tracing of CTCF mutation upon long-term auxin treatment by immunoblot and mass spec analysis.**

**a.** The full CTCF^miniAID3-mClover3^ sequence representing mass spectrometry peptide coverage of the CTCF^miniAID-mClover3^ protein in Clone 3.2 parental and Clones 3.2 and 17.2 resistant cells. Peptides that match the sequence with high confidence are shaded yellow and green.

**b.** Sequence illustration pinpointing the trypsin cleavage breakpoint observed by IP-MS in Clone 17.2 resistant cells. A red asterisk denotes the C > T mutation identified by RNA-seq of resistant cells.

**c and d**. Immunoblot from long-term 5-Ph-IAA culture of Clone 17.2 kept under continuous 1 μM 5-Ph-IAA/ 100 μg/mL Zeocin. Lysates were probed for CTCF and miniAID, and GAPDH was included as a loading control. Two replicates are shown. The auxin-resistant truncated-CTCF appeared around treatment day 18 of rep 1 and treatment day 25 of rep 2.

**Supplementary Fig. 4 CTCF ChIP-seq of CTCF^AID2^ clones expressing either CTCF^WT-HA^ or CTCF^Q666*-HA^**

**a.** Immunoblots of CTCF^AID2^ clones C5, C3.2, and C17.2 with doxycycline-inducible exogenous expression of CTCF^WT-HA^ or CTCF^Q666*-HA^. Cells were treated for 24 hours with 1µM 5-Ph-IAA to remove endogenous CTCF. After 6 hours of treatment, 0.05 µg/mL or 1.0 µg/mL doxycycline was added to CTCF^AID2-WT-HA^ or CTCF^AID2-Q666*-HA^ clones, respectively. The HA and CTCF antibodies were used to assess exogenous CTCF protein expression. GAPDH antibody was used as a loading control.

**b.** Principal component analysis (PCA). PC1 showed 39.3% variance between the CTCF^AID2-WT-HA^ or CTCF^AID2-Q666*-HA^ clones. PC2 showed only 18.8% variance among the replicates.

**c.** Heatmap of CTCF ChIP-seq of CTCF^AID2-WT-HA^ or CTCF^AID2-Q666*-HA^ clones C5, C3.2, and C17.2 showed increased CTCF peak density at 7,727 CTCF binding peaks in CTCF^AID2-WT-HA^ vs CTCF^AID2-Q666*-HA^ clones (log2 FC > 1, FDR < 0.05). Peak density was reduced at only 59 CTCF binding peaks when comparing CTCF^AID2-WT-HA^ vs CTCF^AID2-Q666*-HA^ clones (log2 FC > 1, FDR < 0.05).

**d.** CTCF motif analysis of the 7,727 CTCF binding peaks with increased density in CTCF^AID2-WT-HA^ vs CTCF^AID2-Q666*-HA^ clones showed that the CTCF motifs were most enriched.

**e.** CTCF ChIP seq tracks of CTCF^AID2-WT-HA^ and CTCF^AID2-Q666*-HA^ clones (C3.2 and C17.2) at the *BLCAP* locus showed increased CTCF binding in CTCF^AID2-Q666*-HA^ clones. No variance in binding was observed at most CTCF binding loci, as represented at the *ISYNA1* locus.

**Supplementary Fig. 5 No auxin-inducible degradation resistance observed in human miniAID-RBM5 and MBNL1-miniAID knock-in cell lines.**

**a.** Schematic diagram of the HA mini-AID knock-in design to the N-terminus of *RBM5* in SEM cells. Left-HA and Right-HA designate the homology arms (HA) that flank the targeted knock-in site. Immunoblots for RBM5 and HA showing that ^HA-miniAID^RBM5 remained sensitive to auxin throughout continuous 1 μM 5-Ph-IAA/ 100 μg/mL Zeocin treatment.

**b.** Schematic diagram of the HA mini-AID knock-in design to the C-terminus of *MBNL1* in SEM cells. Immunoblots for MBNL1 and HA showing that MBNL1^HA-miniAID^ remained sensitive to auxin throughout continuous 1 μM 5-Ph-IAA/ 100 μg/mL Zeocin treatment.

**Supplementary Fig. 6 Full characterization of Ctcf-miniAID knock-in mice and primary BCR-ABL B-ALL cells.**

**a.** Successful knock-in was confirmed by genotyping PCR in founder mice. The genotyping PCR primers were combined to detect the 5’ and 3’ junctions of the miniAID. 5’ F (red forward arrow) and R (blue arrow) primers, KI= 478bp; 3’ F (black arrow) and R (red reverse arrow) primers, KI= 448bp; Homology arm primers outside the inserted miniAID sequences (red arrows): WT= 689bp, and KI= 893bp. A wild-type mouse was used as a negative control.

**b.** Successful germline transmission to the F1 progeny was confirmed by genotyping PCR. Primers were designed against the flanking regions of the miniAID cassette (red arrows). Tissues from wild-type and founder knock-in mouse 3F were used as negative and positive controls, respectively.

**c.** Immunoblotting of Ctcf-miniAID expression from tissues, including liver, spleen, and kidney of wild-type, heterozygous, and homozygous knock-in mice. All tissues expressed Ctcf-miniAID fusion protein with a molecular weight 7kDa higher than the wild-type Ctcf protein.

**d.** Simple schematic diagram representing BCR-ABL translocation and sensitivity to second-generation tyrosine kinase inhibitor. The translocation occurs between the ABL gene on chromosome 9 and the BCR gene on chromosome 22. Two translocation fusion proteins are observed: p210, a hallmark of chronic myeloid leukemia (CML), and p185, associated with ALL. TKI, tyrosine kinase inhibitor.

**e.** MTT assay was conducted in parental Ctcf^miniAID/miniAID^ BCR-ABL B-ALL, following a 3-day treatment with Dasatinib in a dosage-dependent manner. K562 was used as a positive control cell line carrying the endogenous translocation of BCR-ABL, and SEM was used as a non-BCR-ABL negative control cell line, replication=3.

**f.** Bright field image demonstrating that Ctcf^miniAID/miniAID^ BCR-ABL B-ALL cells are highly proliferative. Flow cytometry was conducted to show the high expression of B220 and the lack of IgM in the homogenous populations. The eGFP channel indicated the expression of BCR-ABL from MSCV-BCR-ABL-IRES-eGFP.

**g.** Ctcf and miniAID immunoblot of Ctcf^miniAID/miniAID^ BCR-ABL B-ALL cells showed Ctcf degradation after 6-hour 5-Ph-IAA treatment that was restored after auxin removal. Hsp70 was included as a loading control.

**h.** Schematic diagram illustrating CPA assay: Ctcf^miniAID/miniAID^ BCR-ABL B-ALL cells were infected with Cas9 and sgRNA-CFP against the coding exons of *Ctcf*. Flow cytometry analysis of the CFP fluorescence percentage of the cells at day 2, day 5, and day 8 was traced.

**i.** CPA analysis showed time-dependent decreased cell proliferation after Ctcf knock-out. sgNT was included as a negative control, and sgMyc was included as a positive control.

**Supplementary Fig. 7 Establishment of the *in vivo* Ctcf^miniAID/miniAID^ BCR-ABL B-ALL mouse model and characterization of acquired auxin resistance following long-term *in vivo* 5-Ph-IAA treatment.**

**a.** MTT assay showed Ctcf^miniAID/miniAID^ BCR-ABL B-ALL cells after long-term 5-Ph-IAA treatment were still sensitive to Dasatinib, similar to parental cells. SEM cells were included as a negative control.

**b.** Immunoblotting of Ctcf in Ctcf^miniAID/miniAID^ BCR-ABL B-ALL cells treated long-term (day 29) with 5-Ph-IAA showed acquired auxin resistance. HSC70 was included as a loading control.

**c.** Schematic diagram showing *in vivo* model establishment: 100K cells were injected through the tail vein. Weekly peripheral blood was collected for flow analysis until the endpoint.

**d.** Flow analysis of weekly leukemia cell percentage (GFP+ cells) from peripheral blood.

**e.** Image showing spleen size after Ctcf^miniAID/miniAID^ BCR-ABL B-ALL cell injection compared with the negative controls.

**f.** Spleen weight/ body weight of the Ctcf^miniAID/miniAID^ BCR-ABL B-ALL cell-injected mice compared with negative controls.

**g.** Spleen H&E staining of the control and Ctcf^miniAID/miniAID^ BCR-ABL B-ALL cell-injected mice.

**h.** Spleen eGFP staining of the control and Ctcf^miniAID/miniAID^ BCR-ABL B-ALL cell-injected mice.

**i.** After Ctcf^miniAID/miniAID^ BCR-ABL-B-All cells were infected with luciferase-YFP, immunoblotting of Ctcf, miniAID confirmed Ctcf-miniAID fusion protein degradation after 6-hour 5-Ph-IAA treatment before *in vivo* transplantation.

